# Embryonic TGF-β signaling imposes persistent changes to HSPC clonality and inflammatory landscape

**DOI:** 10.64898/2026.07.30.741837

**Authors:** Marta Mastrogiovanni, Anastasia Nizhnik, Joaquín Cantón-Sandoval, Sofia de Oliveira, Teresa V. Bowman

## Abstract

Embryonic hematopoiesis is essential for establishing lifelong blood and immune system function. During development, hematopoietic stem and progenitor cells (HSPCs) acquire intrinsic programs that persist into adulthood and can influence disease susceptibility, yet the molecular signals governing these early-life decisions remain poorly understood. Here, we investigated the role of developmental Transforming Growth Factor-β (TGF-β) signaling in regulating HSPC lineage bias and long-term hematopoietic outcomes. Using the zebrafish model, we found that transient embryonic TGF-β signaling inhibition during the HSPC specification window altered their frequency and migration after emergence from the hemogenic endothelium and movement into the key secondary maturation and expansion niche. Single-cell transcriptomic analysis of embryonic HSPCs revealed repression of migration- and cytoskeleton-associated genes alongside dampened expression of myeloid/macrophage-related genes following ALK5 inhibition. Functionally, early ALK5 blockade reduced macrophage numbers and promoted an M2-like immunosuppressive transcriptional profile. These developmental perturbations produced sustained effects on hematopoietic and immune function into adulthood, including diminished inflammatory gene expression, reduced clonal complexity, and impaired regenerative capacity. Together, our findings identify embryonic TGF-β signaling as a key developmental regulator of HSPC fate and immune programming, with potential implications for immune dysfunction and susceptibility to inflammatory-related disease later in life.

**PAPER HIGHLIGHTS:**

– Transient developmental signaling perturbations reshape hematopoietic trajectories
– Embryogenic TGF-β signaling instructs HSPC lineage priming and macrophage specialization
– Early HSPC programming establishes persistent inflammatory states, clonal diversity, and modifies regenerative capacity

## INTRODUCTION

The blood system is comprised of cells in the lymphoid, myeloid, and erythroid lineages that support organismal health through a multitude of functions such as fighting infections, oxygenating tissues, and repairing injuries. The balanced commitment of hematopoietic stem and progenitor cells (HSPCs) across these lineages is essential for the establishment and lifelong maintenance of blood and immune system homeostasis, and in shaping the functions of both systems. This equilibrium is sustained via the functioning of a pool of HSPCs that combined generate a balanced lineage output. However, on an individual cell level, HSPCs can display lineage bias, which is the preferential generation of a specific hematopoietic lineage more than others. Emerging evidence indicates that these biases are engineered during embryogenesis and can durably influence HSPCs and the number, type, and functional specialization of their mature cell progeny [1–3]. These early life HSPC imprints can also shape predisposing factors for leukemic transformation and/or functional defects later in life [4, 5]. Dissecting the mechanisms that operate during embryonic hematopoiesis is therefore essential to understand how early-life cues instruct later life hematopoietic trajectories and disease risk.

Multiple signaling pathways are implicated in regulating embryonic HSPC emergence and differentiation [6–9]. While many of these developmental cues were characterized for their immediate roles during hematopoietic establishment, far less is known about whether and how transient perturbations of these pathways durably shape postnatal hematopoietic function and disease susceptibility. Indeed, the seeds of hematopoietic disorders can arise from alterations in lineage balance and inflammatory changes, long before overt disease manifests. Therefore, embryonic environmental and signaling perturbations may establish long-term risk-modifying hematopoietic states.

Among the pathways implicated in both developmental hematopoiesis and adult hematopoietic disorders, Transforming Growth Factor-β (TGF-β) signaling represents a key environmental cue. TGF-β has been extensively studied as a regulator of adult HSPC quiescence, lineage bias, and immune responses [10–12]. Also, adult myeloid-biased and lymphoid-biased HSPC subsets display distinct sensitivities to this pathway [11, 13]. Previous studies illustrated that embryonic TGF-β contributes to HSPC specification during the endothelial-to-hematopoietic transition [14, 15], but whether developmental modulation of TGF-β signaling durably shapes adult hematopoiesis remains unknown.

Here, we delved into the developmental role of TGF-β, examining how it regulates HSPC behavior and differentiation during embryogenesis, and if the early-life instructions exert persistent effects on the hematopoietic and immune systems. We show that TGF-β signaling shapes HSPC frequency and migratory dynamics within embryogenic hematopoietic niches and influenced lineage commitment trajectories. Our findings reveal that transient disruption of embryonic TGF-β signaling alters subsequent macrophage output and skews their differentiation toward an immunoregulatory M2-like state. These inflammatory alterations persist into adulthood and are accompanied by changes in hematopoietic clonality, particularly in the myeloid compartment, as well as impaired wound healing responses. Together, our results reveal how transient perturbations during embryonic hematopoiesis can durably reprogram inflammatory competence and clonal architecture later in life, underscoring the importance of gaining deeper insight into embryonic hematopoiesis for improved understanding of later-arising blood and immune diseases.

## RESULTS

### Embryonic inhibition of TGF-β signaling alters HSPC frequency and dynamics within the hematopoietic niche

TGF-β was previously identified among the embryogenic signals shaping HSPC formation but the long term impacts of this perturbation on the hematopoietic system were not explored [14]. Studies perturbing TGF-β signaling in adult mice revealed a selective effect on myeloid-biased HSPC subtype compared to the lymphoid-biased subset [11, 16]. We hypothesized that since TGF-β signaling regulates embryogenic HSPC formation and influences adult HSC lineage preference and immune responses, it could be plausible that this signaling pathway might function to establish these features in the nascent hematopoietic system.

There are numerous TGF-β family ligands connected to HSPC biology [16–20]. In the prior zebrafish study, TGF-β Receptor I/ALK5, which is activated by several TGF-β family ligands important in hematopoiesis, was identified as a key regulator of HSPC emergence [14]. To test our model, we examined the effects of treating zebrafish embryos transiently with the ALK5/TGF-β Receptor inhibitor SB431542 during the critical window of initial HSPC emergence and then examined immediate and long-term consequences of this momentary inhibition (**Figure 1A**). We validated that the TGF-β pathway was inhibited in the larvae by transcriptional analysis of canonical target genes (**Figure S1D**).

**Figure 1:**
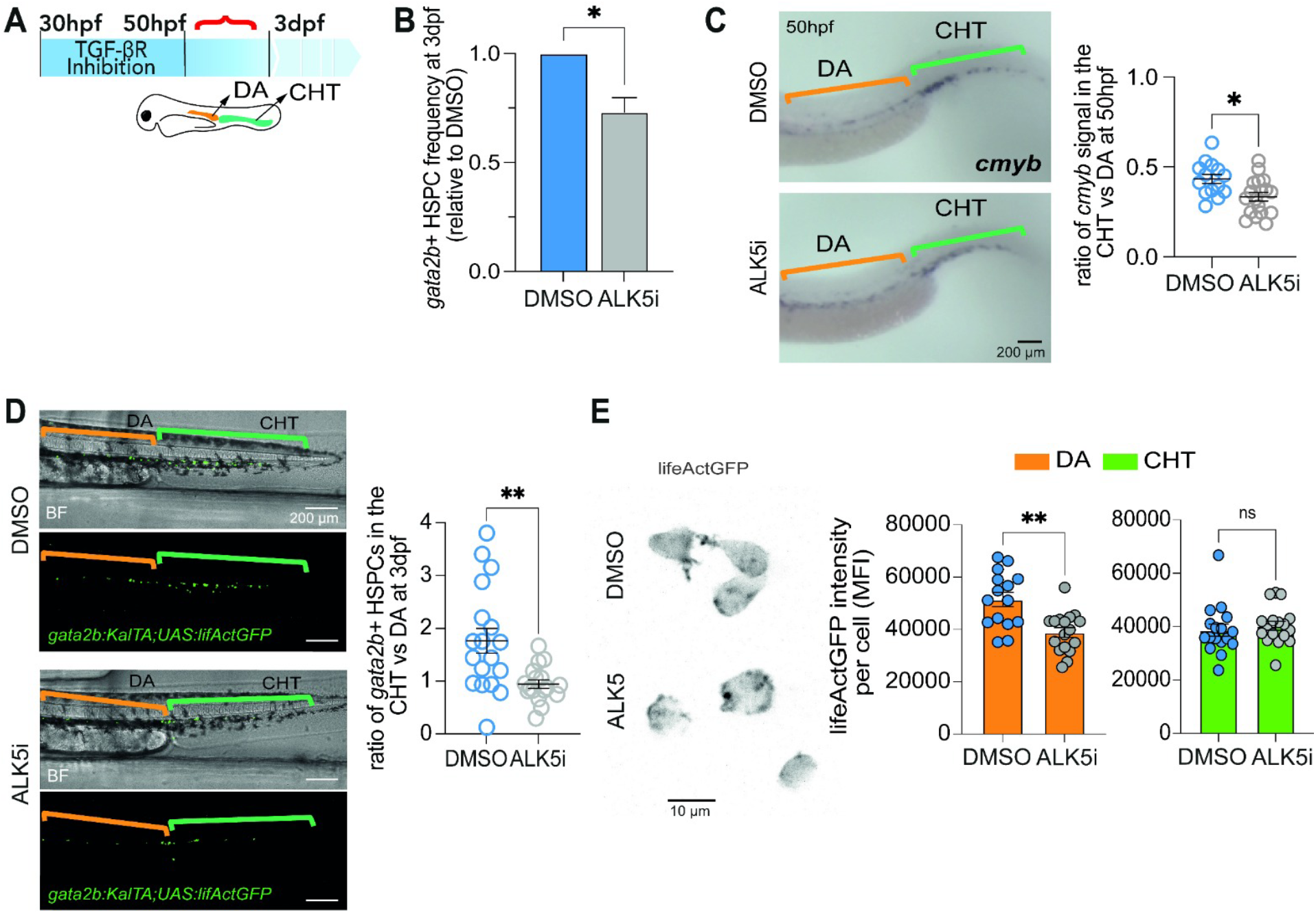
HSPC frequency and localization within hematopoietic niches are altered after TGF-β Receptor inhibition. **A**. Zebrafish embryos were treated between 30 and 50 hpf with the ALK5/TGF-β Receptor inhibitor SB431452 or DMSO vehicle-control. At this time point, HSPCs are emerging from the DA. Analyses were carried out between 50 hpf and 3 dpf (indicated by red bracket), when HSPCs delaminate from the DA and migrate to the CHT. **B**. HSPC frequency in *Tg(gata2b:KalTA;UAS:lifActGFP*) larvae was quantified by flow cytometry at 3 dpf. Bar graph represents mean + SEM of 5 biological replicates. **C**. HSPC localization within the DA and CHT at 50 hpf as assessed by *cmyb* RNA *in situ* hybridization. Left panels show *cmyb*^+^ HSPCs distribution within the DA (orange brackets) and CHT (green brackets). Scale bars, 200µm. Plot on the right shows the relative ratio of HSPCs in the CHT *vs* DA, each dot represents one larva (14-18 total larvae). **D,E**. *Tg(gata2b:KalTA;UAS:lifActGFP*) larvae were imaged at 3 dpf. **D.** Confocal images display the distribution of *gata2b^+^* HSPCs from a representative pair of 3 dpf larvae previously treated with DMSO or ALK5 inhibitor. Brightfield (BF) images are used for the visualization of the two hematopoietic niches of interest, DA (orange brackets) and CHT (green brackets). Scale bars, 200µm, 20x objective. Right graph shows quantification of the relative ratio of HSPCs in the CHT *vs* DA (mean ± SEM). Each dot represents one larva (17 total larvae, 2 biological replicates). **E.** Confocal images of a representative pair of *gata2b^+^*cells from 3 dpf larvae previously treated with DMSO or ALK5 inhibitor. Scale bar 10 µm, 63x objective. Right graphs are the analyses of the Mean Fluorescence Intensity (MFI) of the Filamentous Actin from each larva was measured in the two hematopoietic niches. Graph shows mean + SEM with each dot representing one larva (15-17 total larvae). Statistical differences were calculated by Mann-Whitney unpaired test. *P < 0.05, **P ≤ 0.01.

In zebrafish, HSPCs primarily emerge within the Dorsal Aorta (DA) from approximately one to two days post fertilization (dpf) and subsequently migrate to the caudal hematopoietic tissue (CHT) around 3 dpf where differentiation occurs before colonizing the thymus and the kidney marrow [21–23]. For all of our experiments, we exposed embryos to the ALK5 inhibitor SB431542 from 30-50 hours post fertilization (hpf) to inhibit TGF-β signaling during the time window of HSPC emergence within the DA.

At 3 dpf, ALK5 inhibitor-treated larvae exhibited a reduced total number of *gata2b^+^* HSPCs as measured by flow cytometry of *Tg*(*gata2b:KalTA;UAS:lifActGFP*) animals [24] (**Figure 1B**). Similar results were obtained in *Tg(runx1+23:mCherry*) 3 dpf larvae (**Figure S1A**). To decipher if this represented a decrease within a specific HSPC niche, we investigated HSPC spatial distribution. Immediately after drug treatment at 50 hpf, we observed an altered redistribution of *gata2b^+^*HSPCs between the two principal hematopoietic compartments at this developmental stage: the DA and the CHT (**Figure 1C**). Specifically, ALK5-inhibited embryos displayed preferential accumulation of HSPCs within the DA, accompanied by a reduced presence in the CHT, as assessed by *cmyb* whole mount *in situ* hybridization (**Figure 1C**). Quantification of *gata2b*^+^ HSPCs by live imaging confirmed that the DA-to-CHT ratio remained skewed in ALK5-inhibited larvae at 3 dpf (**Figure 1D**). These findings suggest defective colonization of the CHT niche and reveal both frequency and distribution defects at this critical embryogenic developmental stage upon diminished TGF-β signaling.

HSPCs rely on their migratory dynamics both to transition between hematopoietic niches and to establish the appropriate spatial distribution within each niche, enabling proper engagement with niche cells, neighboring cell populations, and specific molecule gradients [25–27]. Given that cell migration implies remodeling of the actin cytoskeleton, including the polymerization of globular actin into Filamentous-actin (F-actin) [28–30], we first investigated the F-actin network in LifAct:GFP-expressing *gata2b^+^*cells. ALK5 inhibition resulted in clear changes in F-actin polymerization (**Figure 1E**), specifically decreasing F-actin levels in cells residing within the DA with no significant difference noted in the CHT (**Figure 1E**, right panel). This finding strongly correlates with their accumulation in this compartment and suggests impaired delamination and/or relocalization from the DA to the CHT.

To determine whether the defective CHT colonization and altered F-actin organization observed following ALK5 inhibition were associated with changes in HSPC migratory behavior, we tracked *gata2b^+^*cell behaviors within the DA and the CHT right after drug treatment, between 50 and 58hpf (**Figure 2, Supplemental Videos 1,2**). Following treatment with DMSO or the ALK5 inhibitor, we performed live imaging of GFP^+^ cells from *Tg*(*gata2b:KalTA;UAS:lifeActGFP*) larvae. To specifically characterize HSPC dynamics within these hematopoietic niches, we focused imaging on the DA and CHT regions at high temporal resolution (2-minute intervals), enabling precise tracking of individual HSPC migratory behaviors over time. Thereafter, we extracted multiple kinetic parameters describing single cell dynamics over time in each anatomical compartment, enabling the construction of behavioral descriptors as previously described [31]. This approach allowed us to identify HSPC behavioral clusters within the DA vs CHT hematopoietic niches and under DMSO and ALK5-inhibited conditions. Based on their migration behavior, we identified 8 and 16 distinct clusters in the DA and CHT, respectively (**Figure 2A-D**).

**Figure 2:**
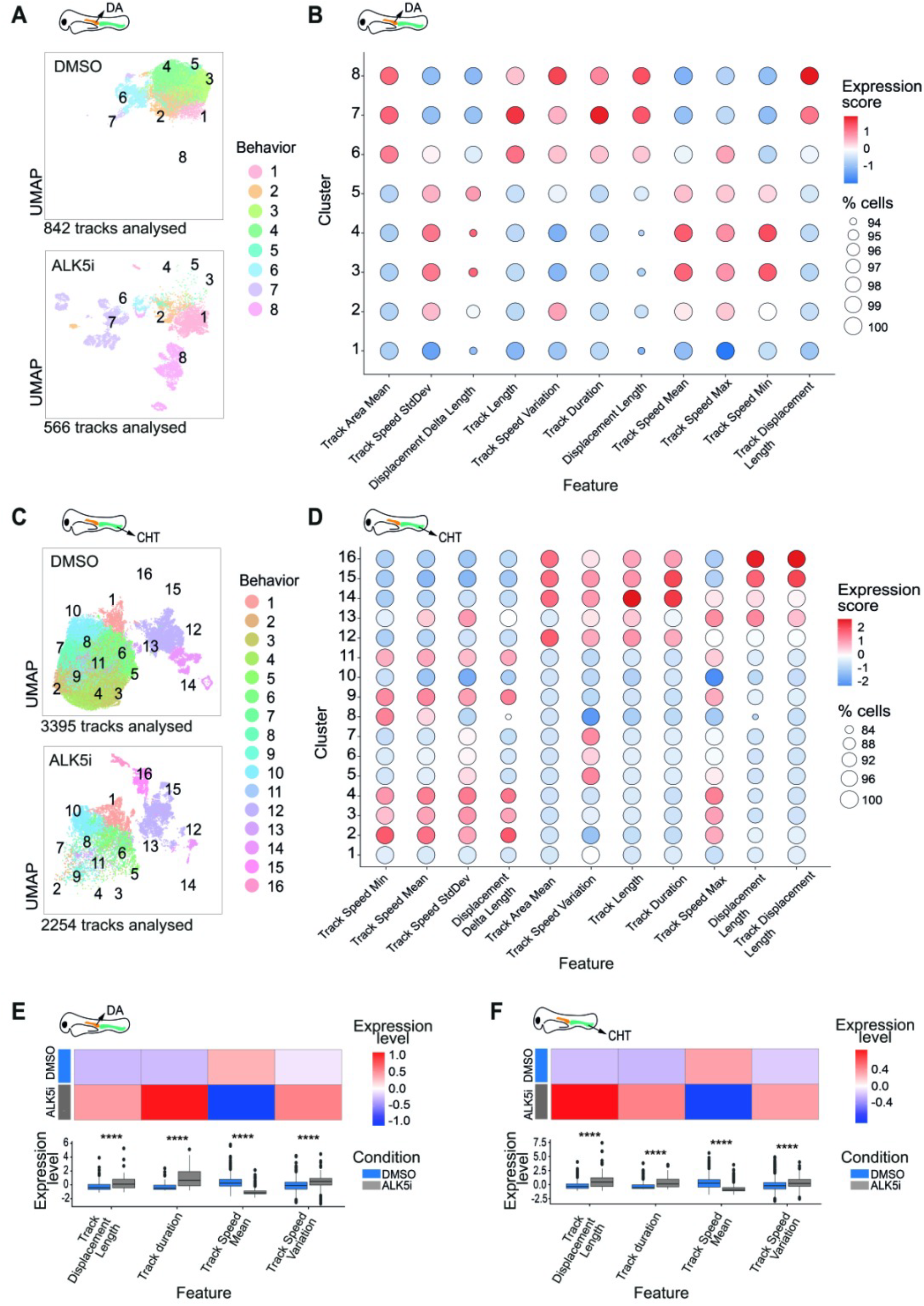
HSPC migratory dynamics within the hematopoietic niches are altered after TGF-β Receptor inhibition. Zebrafish embryos were treated between 30 and 50 hpf with the ALK5 inhibitor SB431452 or DMSO vehicle-control. HSPC cell migration dynamics within DA and CHT were assessed by live imaging of 2-3 *Tg(gata2b:KalTA;UAS:lifActGFP*) embryos per condition from 2 independent experiments. Migration was recorded for 8 hours starting from 50 hpf using spinning disk confocal, 10x objective. **Supplemental Videos 1** and **2** display the migration of HSPCs in, DA (1) and CHT (2) from a representative pair of DMSO and ALK5 inhibitor-treated embryos. **A,C.** UMAP representations of HSPC single-cell tracking data from DMSO and ALK5 inhibitor-treated embryos in DA (**A**) and CHT (**C**). Tracks were analyzed with each track corresponding to a single HSPC, and clustered based on HSPC features of motion, resulting in 8 and 16 behavioral clusters, respectively in DA (**A**) and CHT (**C**). UMAPs are split by condition (DMSO-top *vs* ALK5 inhibitor-bottom) and colored by behavioral cluster identity. **B,D.** Dot plots showing the main kinetics parameters across the different behavioral clusters. Dot size represents the percentage of cells within each cluster with measurable values for a given feature, while color indicates the z-scored normalized value of the parameter, with red and blue representing, respectively, cluster enrichment and depletion of a specific behavior relative to the mean expression across all cells. DA data shown in (**B**) and CHT data shown in (**D**). **E,F.** Heatmaps combined with boxplots showing statistical differences in migratory features of HSPCs between DMSO and ALK5 inhibitor-treated embryos in the DA (**E**) and CHT (**F**). Heatmaps (top) display z-scored feature values (red: higher than average feature values, blue: lower than average values, white: centered at 0, corresponding to the average) across conditions. Boxplots (bottom) display the distribution of z-scored values for each feature in DMSO (blue) and ALK5 inhibitor-treated (gray) conditions. Boxplots indicate median and interquartile range of single HSPC tracks analyses. Statistical significance was assessed using a two-tailed Mann–Whitney test with multiple testing correction (Benjamini–Hochberg). ****P ≤ 0.0001.

In the DA, ALK5 inhibition induced a marked shift in HSPC migratory behavior (**Figure 2A**). While DMSO-treated cells were predominantly distributed across clusters 3–5, ALK5-inhibited HSPCs became enriched in clusters 7 and 8, revealing a substantial reorganization of behavioral states within the niche (**Figure 2A**). Notably, examination of the migration parameters defining these states showed that ALK5-inhibited HSPCs displayed reduced track speed together with increased displacement length compared with control cells (**Figure 2B,E**). In contrast, the behavioral states enriched in DMSO-treated embryos were characterized by higher migratory speed and lower displacement length (**Figure 2B,E**). This finding suggests that embryonic TGF-β signaling regulates distinct HSPC migratory programs consistent with prolonged searching within the DA, potentially contributing to their retention within the site of emergence and reduced colonization of the CHT.

The CHT was characterized by a greater heterogeneity of migratory behaviors than in the DA (**Figure 2C**), consistent with prior studies [32, 33]. Although the overall distribution of cells across clusters was less dramatically altered by ALK5 inhibition than in the DA, ALK5-treated HSPCs retained key migratory features observed at the site of emergence, including reduced track speed and increased displacement length (**Figure 2D,F**). The persistence of these behavioral alterations across both hematopoietic niches suggests that embryonic TGF-β signaling regulates intrinsic migratory programs that accompany HSPCs during niche transition rather than exerting purely niche-specific effects.

Collectively, these findings establish a novel framework for characterizing embryonic HSPCs based on dynamic behavioral properties. By integrating the assessment of spatial redistribution within hematopoietic niches and quantitative migration descriptors, we reveal a previously unappreciated role for TGF-β signaling in regulating embryonic HSPC dynamics during niche colonization.

### Embryonic TGF-β signaling alters the HSPC transcriptome and is a key determinant of early HSPC myeloid lineage bias and macrophage differentiation

In attempts to gain additional insight into regulatory mechanisms influencing HSPC dynamic behaviors within the hematopoietic niches, we interrogated whether DMSO vs ALK5-inhibited HSPCs presented differences in their transcriptional programs. To this end, we did single-cell RNA sequencing of *gata2b^+^* cells isolated from 3dpf *Tg(gata2b:KalTA;UAS:lifActGFP*) larvae treated earlier with DMSO or ALK5 inhibitor (**Figure S1B**). We identified 3 clusters of HSPCs defined by expression of the HSPC marker *cmyb*. Although the ALK5 condition displayed fewer total HSPCs consistent with our flow cytometric findings (**Figure 1B and S1A**), the distribution of

HSPC clusters was similar between DMSO and ALK5 inhibited conditions (**Figure S1C**). Cluster 0 was characterized by the expression of lymphoid progenitor markers such as *ikzf2* and *runx3*, a proliferative Cluster 1 expressing markers such as *mki67* and *cdk1*, and a HSC-like Cluster 2 expressing *drl* and *apln* genes (**Figure 3A-C and Supplementary File 1**). Focusing on the potential functional differences between the HSPCs by condition, we next interrogated the differentially expressed genes between DMSO and ALK5-inhibited samples for enrichment in different pathways using GSEA GO. Consistent with the altered HSPC migratory behavior observed in ALK5-inhibited embryos (**Figure 1,2**), our scRNA-seq result revealed significant differences in multiple pathways associated with cell migration (**Figure 3D**). Notably, multiple pathways involved in cell adhesion and motility were significantly enriched in the DMSO condition compared to ALK5-inhibited HSPCs (**Figure 3D**). Specific genes in these pathways included integrins such as *icam3* and *itgb4*, as well as chemokines such as *cxcl19* (**Figure 3E**). DMSO-treated HSPCs also showed higher expression of key regulators of actin filament organization and polymerization, including *cdc42bpb* and *rac1a* (**Figure 3D,E**). Together, these transcriptional differences align with the observed migration defects and F-actin phenotypic alterations (**Figures 1,2**), supporting a coherent link between disrupted motility and cytoskeletal gene programs and altered HSPC cellular dynamics.

**Figure 3:**
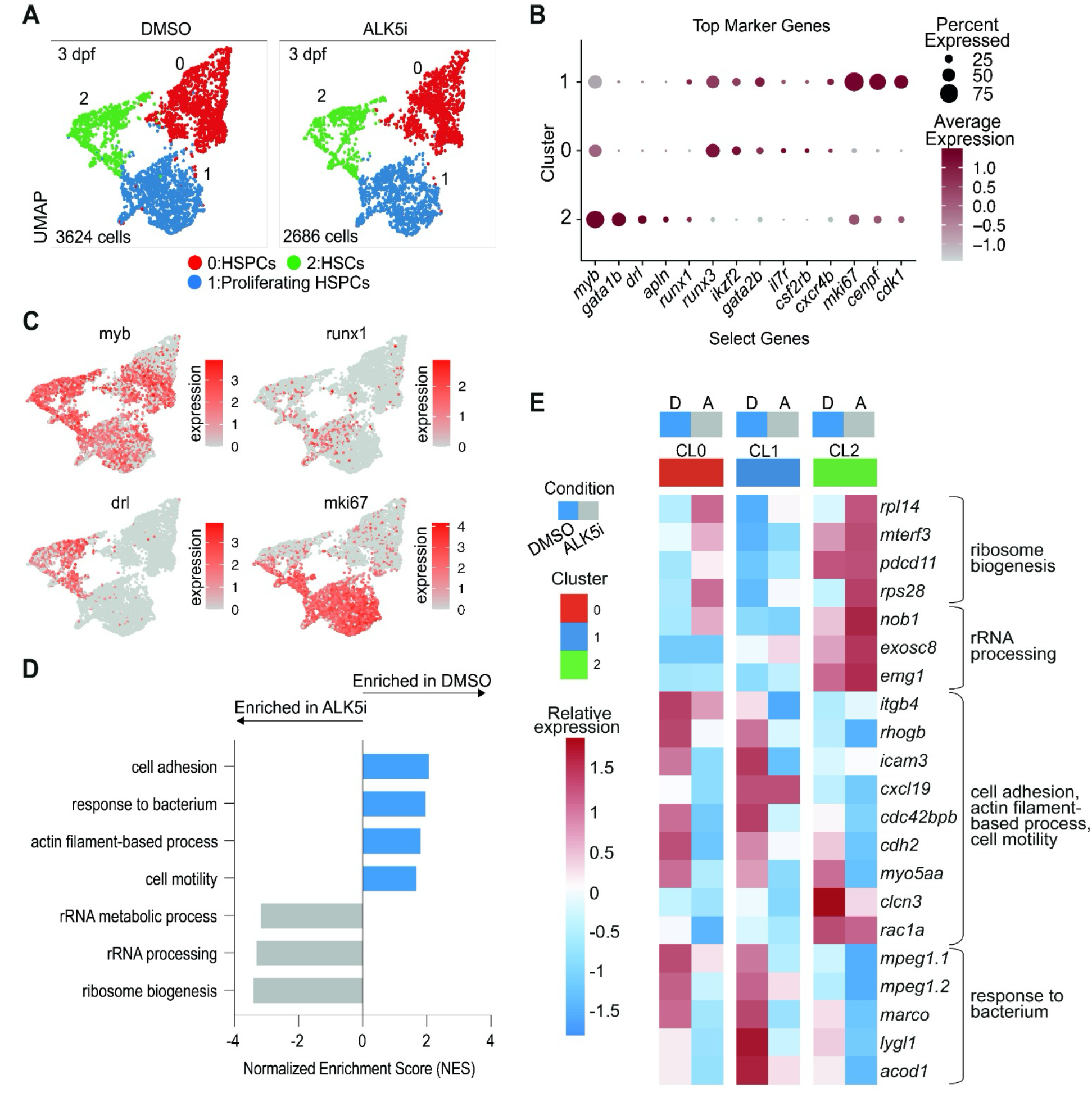
Embryonic TGF-β Receptor inhibition alters larval HSPC transcriptional signature. **A.** UMAP visualization of *gata2b^+^* HSPCs from 3 dpf *Tg(gata2b:KalTA;UAS:lifActGFP*) larvae treated with DMSO or ALK5 inhibitor from 30-50 hpf. Distinct cell clusters are denoted in different colors and were determined by differential gene expression (**Supplementary File 1**). **B.** Dot plot showing selected top markers differentially expressed across the HSPC clusters. Dot size indicates the proportion of cells within each cluster expressing a given marker and color indicates the mean expression level in that cluster. Full list of top marker genes is in **Supplementary File 1**. **C.** Feature plot displaying the expression of representative HSPC genes including *myb*, *runx*, *drl*, and the proliferation marker gene *mki67*. Red color scale shows relative gene expression enrichment. **D.** GSEAGO pathway analysis comparing HSPCs from ALK5 inhibitor-treated *vs* DMSO conditions. NES (normalized enrichment score) values indicating changes in genes involved in the regulation of the listed biological processes in *gata2b^+^* HSPCs from the three clusters together. The full list of GSE pathways and genes is in **Supplementary File 1**. **E.** Heatmap showing selected differentially expressed gene profiles across the different clusters and conditions. Each row represents a unique gene, which belongs to the functional pathways displayed in D. Each column represents the DMSO vs ALK5 inhibited conditions in each cluster. Relative gene expression levels are signified by different color scale (red=higher and blue=lower).

Proper HSPC localization and migration within the CHT is a crucial step underlying cell fate [21, 32, 34]. As we noted differences in CHT seeding after embryonic ALK5 inhibition (**Figures 1,2**), we investigated if HSPC lineage priming could be defective. Pathway analysis of scRNA-seq data from 3dpf *gata2b^+^* cells revealed that the genes downregulated in the ALK5-inhibited group compared to the DMSO sample were associated with the “response to bacterium” pathway (**Figure 3D**). Notably, these genes largely corresponded to myeloid and macrophage-associated genes, such as *acod1*, *marco, lygl1* and *mpeg1.1* (**Figure 3E and Supplementary File 1**). In contrast, genes expressed higher in the HSPCs from ALK5-inhibited embryos were enriched for factors involved in ribosome biogenesis, including *rpl14* and *pdcd11,* similar to prior observations [35, 36] (**Figure 3D,E and Supplementary File 1**). This transcriptional profile is consistent with a less differentiated or more immature state, supporting the notion of reduced differentiation potential.

To further investigate potential shifts in myeloid bias, we examined the expression of canonical myeloid marker genes in the HSPC scRNA-seq data (**Figure 4A**). Our analyses showed that macrophage signature genes were downregulated in the HSPCs after ALK5 inhibition (**Figure 4A,B**). In addition to those identified by pathway enrichment (**Figure 3D**), other macrophage signature genes such as *mfap4.1* and *csfr1a* were also downregulated (**Figure 4A,B**) upon ALK5 inhibition. Conversely, HSC markers such as *gata2b* and *myb* were trending up in Clusters 0 and 1 in ALK5-inhibited cells, reinforcing a shift toward a less differentiated state. Expression of erythroid and neutrophil-associated genes were also somewhat diminished in HSPCs upon ALK5 inhibition (**Figure 4A and Figure S1E**).

**Figure 4:**
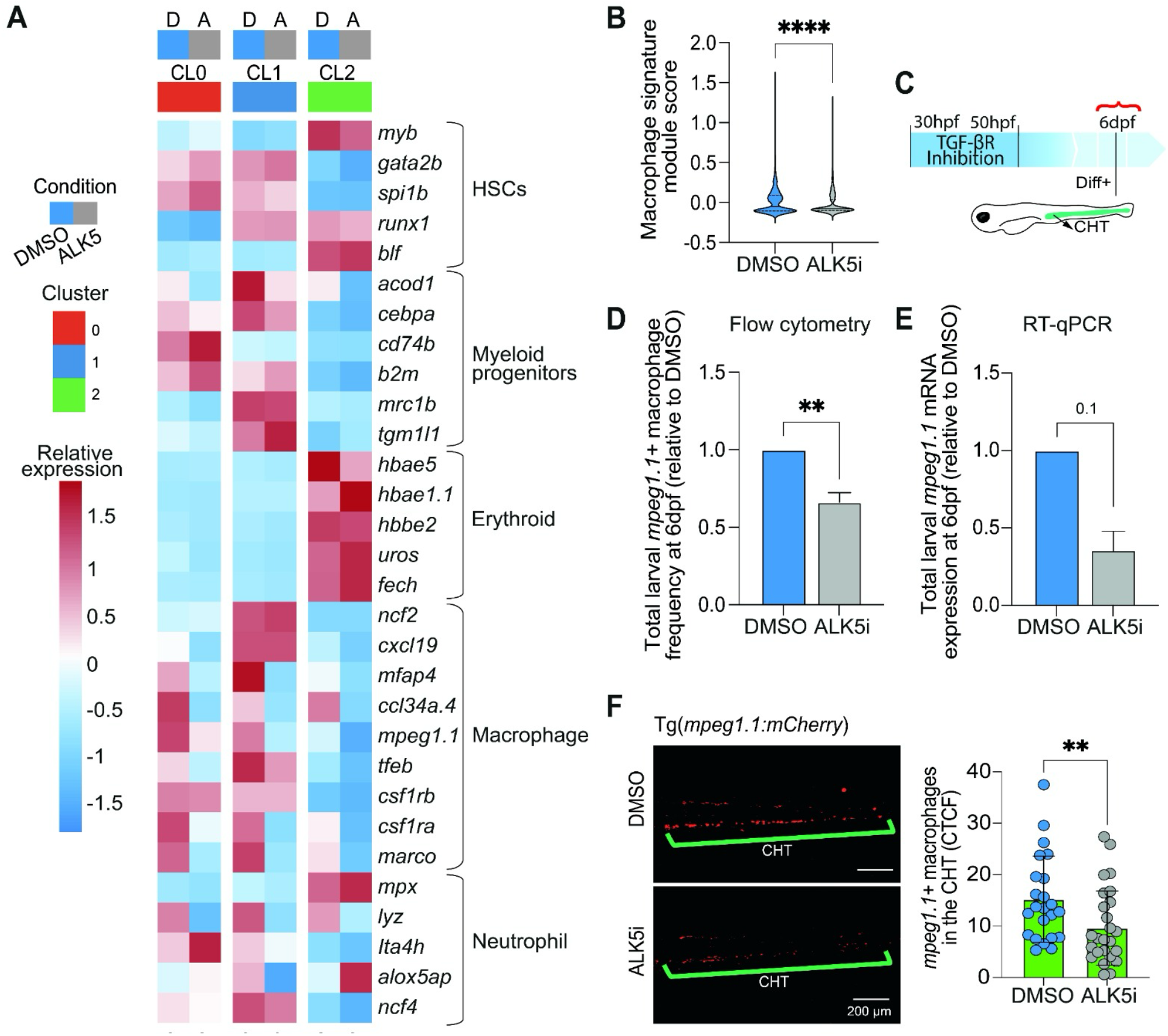
Embryonic TGF-β Receptor inhibition skews HSPC lineage output. **A.** Heatmap showing the relative expression of lineage-specific signature genes across hematopoietic populations in *gata2b^+^* HSPCs from 3 dpf *Tg(gata2b:KalTA;UAS:lifeActGFP*) larvae treated with DMSO or ALK5 inhibitor. Rows are individual genes and columns represent DMSO and ALK5 inhibitor-treated conditions within each identified cluster. Genes are grouped according to hematopoietic lineages, including HSCs, myeloid progenitors, erythroid cells, macrophages, and neutrophils. Color scale indicates relative gene expression levels. **B.** Quantification of macrophage-associated gene module scores in DMSO and ALK5 inhibitor conditions across all clusters. List of genes used for the module score is in **Supplementary File 1**. Statistical differences were calculated by Wilcoxon test. ****P ≤ 0.0001. **C.** *Tg(mpeg1.1:mCherry*) embryos were treated from 30-50 hpf with the ALK5 inhibitor SB431452 and then the number and distribution of *mpeg1.1^+^*cells was assessed at 6 dpf, when differentiated mature populations (Diff+) are abundant in the CHT. **D.** Total frequency of *mpeg1.1:mCherry^+^* macrophages at 6dpf was measured by flow cytometry. Bar graph represents mean + SEM of 6 biological replicates. Values shown are normalized to the DMSO control per experiment. Statistical differences were calculated by Mann-Whitney unpaired test. **P < 0.01. **E.** RT-qPCR quantification of *mpeg1.1* mRNA expression in whole larvae at 6 dpf. Values were normalized to *rps11* and expressed as fold change relative to DMSO. Bar graphs displaying mean + SEM of 3 independent experiments. Statistical differences were calculated by Mann-Whitney unpaired test. **F.** The number of *mpeg1.1:mCherry^+^*macrophages within the CHT at 6 dpf was quantified by fluorescent microscopy and Corrected Total Cell Fluorescence (CTCF). Left panels display the CHT of two representative larvae: DMSO (top) and ALK5 inhibited (bottom). Right bar graph represents mean ± SD of 2 independent experiments. Each dot represents a larva (>12 per replicate). Statistical differences were calculated by Mann-Whitney unpaired test. **P < 0.01.

We next assessed whether the decreased lineage specific transcriptional signatures in ALK5-inhibited embryos translated into decreases in the number of differentiated blood cells at a later larval stage. At 6dpf, embryonic ALK5-inhibited larvae displayed a significant reduction in the total frequency of *mpeg1.1^+^*cells in whole larvae as detected by flow cytometry and of *mpeg1.1* RNA expression as detected by RT-qPCR (**Figure 4D,E**). There was also a pronounced decrease of *mpeg1.1^+^* cells within the CHT of ALK5-inhibited larvae, as assessed by microscopy (**Figure 4F**). In contrast, other differentiated myeloid-derived populations, such as neutrophils and erythrocytes, were not significantly affected (**Figure S1F,G**). Altogether, these findings indicate a selective impairment in macrophage lineage priming following TGF-β inhibition.

### Embryonic TGF-β inhibition promotes an M2-like anti-inflammatory phenotype

Macrophages are key immune cells that acquire distinct pro-inflammatory (M1-like) or anti-inflammatory (M2-like) functional features during differentiation, giving rise to heterogeneous populations with specialized inflammatory and tissue-associated roles. Prior studies suggest that TGF-β can directly impact adult macrophage polarization and function [37–40], but how signaling perturbation in embryonic HSPCs affects macrophage progeny is unknown. To characterize how embryonic TGF-β signaling modulation during HSPC specification influences macrophage differentiation, we performed single-cell transcriptional profiling of *mpeg1.1:mCherry^+^* macrophages at 6 dpf (**Figure 5 and Figure S2**). We identified 4 distinct macrophage clusters, including proliferating and differentiated populations (**Figure 5A,B**). Consistent with our previous observations of reduced macrophage number (**Figure 4D-F**), ALK5-inhibited larvae displayed an overall decrease in macrophage abundance compared to controls, while the relative distribution of cells across clusters remained unchanged (**Figure S2B**). Clusters 0 and 1 comprised the majority of the *mpeg1.1*^+^ population (about 80% of all cells), whereas clusters 2 and 3 represented smaller subsets (<10% each; **Figure S2B**). Marker gene analyses indicated that clusters 2 and 3 transcriptionally resembled clusters 0 and 1, indicating related but less abundant states (**Figure 5B**).

**Figure 5:**
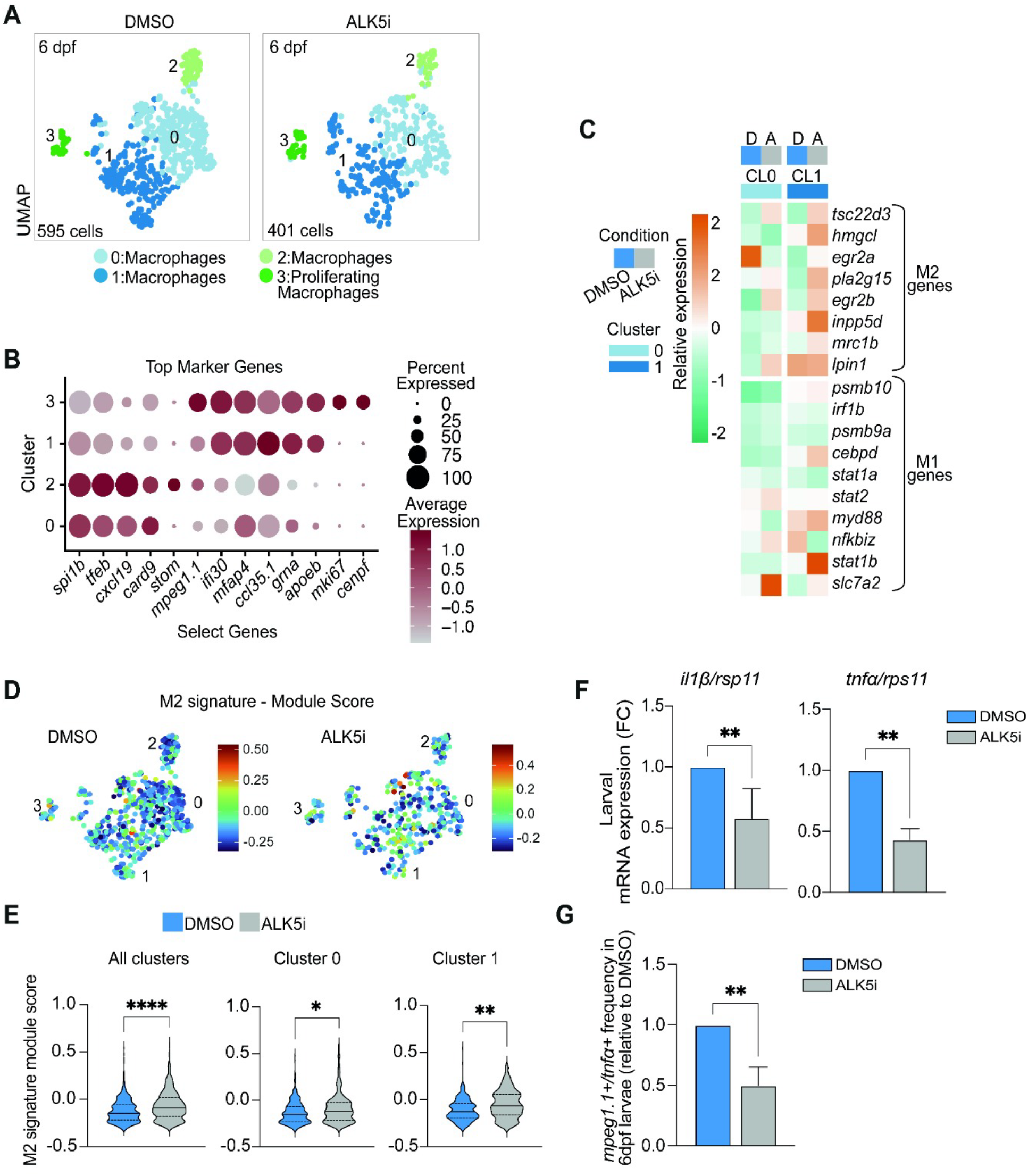
Single cell transcriptomics reveals diverse larval macrophage subsets and different macrophage polarization states upon embryonic TGF-β Receptor inhibition. **A.** UMAP plot displays 4 macrophage cell type clusters in 6 dpf larvae previously treated with DMSO or ALK5 inhibitor from 30-50 hpf: macrophages (main clusters 0,1 and cluster 2) and proliferating macrophages (3). **B.** Dot plots showing expression of a select subset of top marker genes used to designate cluster identity assignments. Dot size represents the percentage of cells expressing each gene with expression level indicated as color intensity. Full list of top marker genes is in **Supplementary File 2**. **C.** Heatmap of M1/M2 polarization-associated genes across the DMSO and ALK5-inhibition conditions in the two main clusters (0 and 1). Genes are grouped into M1-like and M2-like signatures. **D,E.** Module scores for M2-associated genes in DMSO and ALK5 inhibited conditions with quantification across all clusters and within clusters 0 and 1. Full lists of signature genes are listed in **Supplementary File 2**. Color in the feature plots indicates the M2 macrophage module score, where positive values denote enrichment and negative values denote depletion of the M2-macrophage transcriptional program. Statistical differences were calculated by Wilcoxon test. *P < 0.05, **P < 0.01, ****P < 0.0001. **F.** RNA expression of two pro-inflammatory cytokines, *il1β* and *tnfα*, was measured by RT-qPCR in 6 dpf larvae treated previously with DMSO or ALK5 inhibitor. Values were normalized to *rps11* and expressed as fold change (FC) (2 independent experiments). **G**. Single cell suspension from *Tg(tnfα:egfp;mpeg1.1:mCherry)* 6 dpf larvae was processed for flow cytometry. Dead cells were excluded using Draq7 staining. The percentage of *mpeg1.1:mCherry* and *tnfα:eGFP* double positive cells was quantified in 3 independent experiments. Values were normalized to the DMSO and expressed as fold change (FC). Statistical differences were calculated by Mann-Whitney unpaired test. **P < 0.01.

Further analyses of the average gene expression of the two main macrophage clusters suggested a shift in macrophage polarization after early-life ALK5 inhibition. We observed an increase in expression of M2-associated genes alongside reduced M1-associated genes (**Figure 5C**). These included the decreased expression of pro-inflammatory transcription factors, including *nfkbiz,* and the upregulation of immunosuppressive mediators, such as *tsc22d3* (**Figure 5C**). Consistently, module scoring demonstrated an enrichment of M2-like transcriptional programs but not M1-like signatures in all larval macrophage clusters from embryonic ALK5-inhibited animals (**Figure 5D,E and Figure S2C,D**). These findings were further supported by a reduction in pro-inflammatory gene expression and by a decreased proportion of inflammatory *tnfα*^+^ *mpeg1.1*^+^ macrophages, as determined by RT-qPCR and flow cytometry, respectively (**Figure 5F,G**). Together, these data indicate that disruption of embryonic TGF-β signaling not only reduces macrophage output but also skews macrophage differentiation toward an immunoregulatory, M2-like state, supporting a model in which early HSPC programming directly influences downstream myeloid cell identity and inflammatory function.

### Inflammatory defects persist long term and are accompanied by clonal skewing and aberrant wound repair

A primary goal of our study was to determine whether, and to what extent, altered embryonic signaling impacts adult hematopoiesis. Our data show that inhibition of TGF-β signaling during embryogenesis compromises the HSPC compartment resulting in macrophage alterations in both frequency and polarization states. These changes are consistent with embryonic ALK5 inhibition during HSPC emergence promoting a more anti-inflammatory state (**Figure 5C-G**). We therefore assessed whether these alterations are maintained during adulthood. Adult zebrafish treated during embryogenesis with the ALK5 inhibitor exhibited reduced expression of pro-inflammatory cytokines in the whole kidney marrow (WKM), accompanied by reduced *mpeg1.1* cell marker expression (**Figure 6A,B**). Consistently, the proportion of inflammatory *tnfα*^+^ *mpeg1.1^+^* cells were decreased (**Figure 6C**).

**Figure 6:**
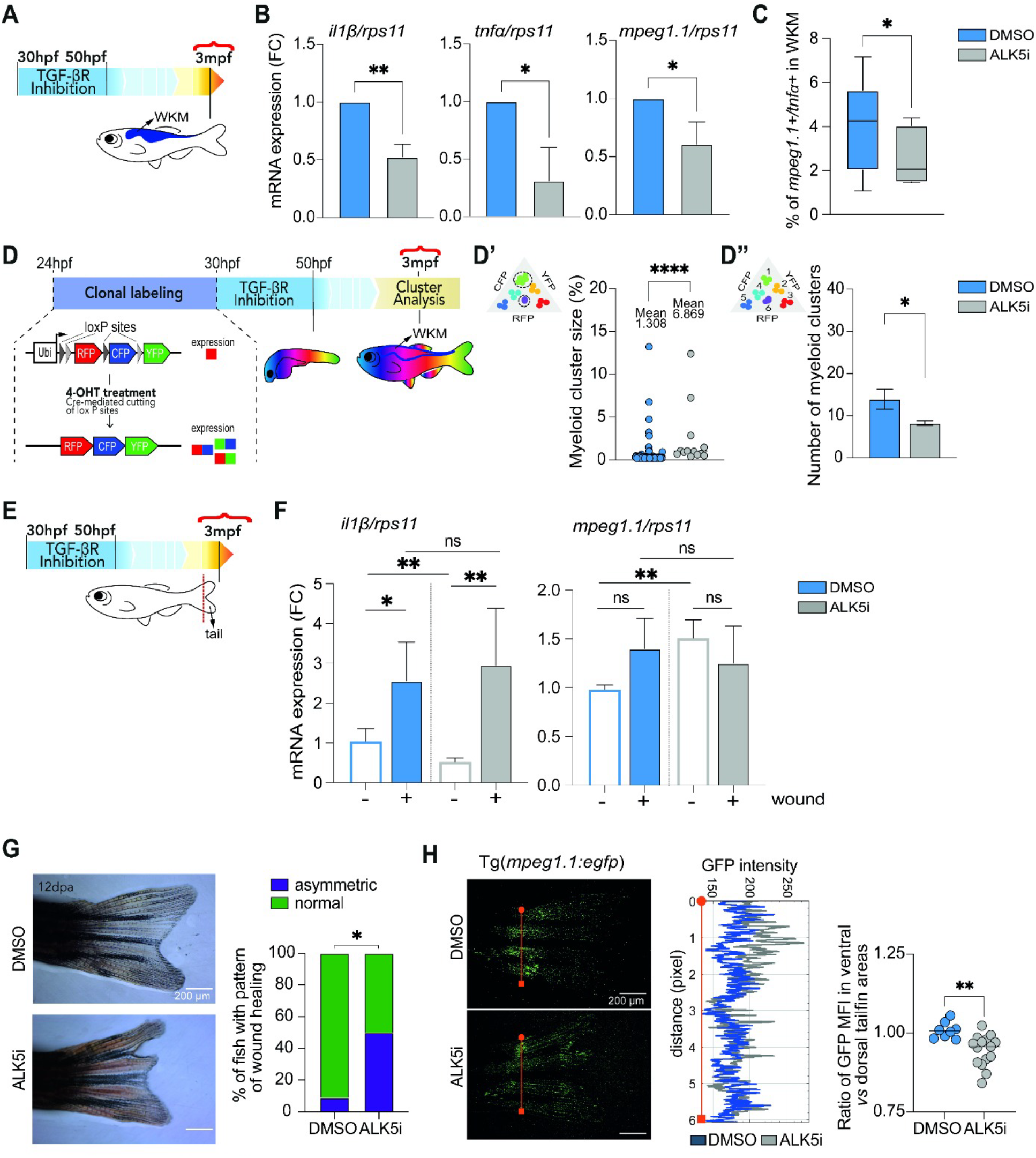
Embryonic inhibition of TGF-β signaling has long term effects on inflammation, clonality and wound response. **A.** Zebrafish embryos were treated with DMSO or ALK5 inhibitor at 30-50hpf and grown to adulthood (older than 3 month post fertilization (mpf)). Experimental schematic showing that whole kidney marrow (WKM) was isolated from adult fish and used for the analyses shown in B and C. **B.** RNA expression of *il-1β*, *tnfα* and *mpeg1.1* was measured by RT-qPCR in adult WKMs from 4 independent experiments. Values were normalized to *rps11* and expressed as fold change (FC). Statistical differences were calculated by Mann-Whitney unpaired test. *P < 0.05, **P < 0.01. **C.** The percentage of *mpeg1.1:mCherry* and *tnfα:eGFP* double positive cells in the WKM from adult *Tg(tnfα:eGFP;mpeg1.1:mCherry*) fish was quantified by flow cytometry in 3 independent experiments, N= 4-6 animals per group. Statistical differences were calculated by Mann-Whitney unpaired test. *P < 0.05. **D.** The Zebrabow transgene encodes for three fluorophores, Cyan Fluorescent Protein (CFP), Yellow Fluorescent Protein (YFP), and Red Fluorescent Protein (RFP) in tandem and with multiple lox-sites. The Zebrabow expressing animals contain up to 40 copies of this transgene. In the inactivated state, all the cells express the RFP fluorophore. Upon 4-OHT-induction, *drl-*driven CreERT2 stochastically excises lox-flanked segments of the Zebrabow transgene, resulting in different levels of the CFP, YFP and RFP, creating a unique “barcode” color for every cell and its progeny. *Tg(ubi:Zebrabow-M;drl:creERT2*) embryos were treated with 4-OHT from 24-30 hpf. They were then treated with DMSO or the ALK5 inhibitor from 30-50 hpf and grown to adulthood. WKM was dissected and myeloid cells defined by FSC and SSC characteristics were analyzed for fluorescent protein expression. Flow cytometry data were clustered using a custom MATLAB graphical tool developed by Avagyan et al., 2021 [41]. Cluster size, which informs on the clonal expansion of the same HSPC clone (D’), and cluster number, which informs on clonal diversity within the HSPC pool (D’’) were analyzed. Two independent experiments, N=16-20 animals per group. Statistical differences were calculated by Mann-Whitney unpaired test. *P < 0.05, ****P < 0.0001. **E**. Tailfin regeneration experiments were performed on adult zebrafish treated embryonically with DMSO or the ALK5 inhibitor. Two independent experiments, N=11-16 animals per group. **F.** After imaging on 0 and 12 days post amputation (dpa), tailfins were resected and used for RT-qPCR quantification. RNA expression of *mpeg1.1* and *il-1β* was measured by RT-qPCR in unwounded (-) and wounded (+) conditions (n=5). Values were normalized to *rps11* and expressed as fold change (FC). Statistical differences were calculated by Mann-Whitney unpaired test. *P < 0.05, **P < 0.01, ns, not significant. **G.** Representative images of 12 dpa caudal fin morphology in adult fish treated with DMSO (top) *vs* ALK5 inhibitor (bottom) during embryonic development. Bar graph depicts percentage of animals with normal *vs* asymmetric fin regeneration patterns. Scale bar, 200 μm. Statistical differences were calculated by Fisher’s Exact Test *P < 0.05. **H.** Representative images of *mpeg1.1^+^* cells in 12 dpa fins from *Tg(mpeg1.1:eGFP*) fish: DMSO (top) and ALK5 inhibitor (bottom). Plot profiling the spatial GFP intensity distribution from the tailfin from dorsal (circle) to the ventral (square) side. Quantification of the ratio of GFP mean fluorescence intensity (MFI) in ventral *vs* dorsal tailfin regions. Each dot represents one fish (2 experiments). Statistical difference was calculated by Mann-Whitney unpaired test. **P < 0.01.

Previous findings demonstrated that imbalanced pro-*vs* anti-inflammatory gene expression influences the clonal fitness of myeloid populations in the WKM, with relevance to clonal hematopoiesis driven by acquired somatic mutations [41]. Given that embryonic modulation of TGF-β signaling perturbed inflammatory gene expression, we next assessed whether it leads to changes in adult hematopoietic cell clonality. To investigate the WKM clonal architecture, we used a hematopoietic-specific multi-color genetic barcoding system (aka Zebrabow) that consists of endogenous fluorescent protein labeling of HSPC clones in zebrafish [41, 42]. To evaluate HSPC clonality, *Tg(Zebrabow-M;drl:creERT2*) embryos were treated with 4-OHT from 24-30hpf, exposed to either DMSO or the ALK5 inhibitor during the main embryonic HSPC specification window (30-50hpf), and then raised to adulthood (>3 months) for clonal analysis of myeloid, lymphoid, and precursor populations as previously described [42] (**Figure 6D, S3A**). Our findings show that ALK5-inhibited animals exhibited a significant increase in myeloid clone size (**Figure 6D’**) accompanied by a reduction in total clone number (**Figure 6D’’**), indicative of decreased clonal diversity and enhanced clonal selection. We also found that the lymphoid and precursor populations showed more modest changes, with increased clone size but no substantial reduction in clone number (**Figure S3B**). Together, these data demonstrate that embryonic TGF-β signaling shapes long-term inflammatory signatures and hematopoietic clonal architecture in the marrow. Its transient disruption during embryogenesis leads to persistent reduction of pro-inflammatory states and promotes myeloid-biased clonal skewing.

Beyond their role in hematopoietic tissues, macrophages are essential regulators of inflammatory homeostasis in peripheral tissues, where, for example, they orchestrate immune responses during tissue regeneration and repair [43, 44]. Regeneration following injury requires a tightly coordinated inflammatory program in which macrophages sequentially transition through distinct inflammation-functional states, initially promoting inflammatory activation, then clearing cellular debris, and ultimately supporting tissue remodeling and inflammatory resolution [45–47]. Given the persistent anti-inflammatory phenotype observed in hematopoietic tissues following embryonic ALK5 inhibition, we next asked whether macrophage-dependent inflammatory responses were also altered in peripheral tissues.

To investigate inflammatory signals associated with macrophage function in peripheral tissues, we analyzed expression of the pro-inflammatory cytokine *il1β* in tailfins from adult zebrafish exposed during embryogenesis to either DMSO or the ALK5 inhibitor during the window of HSPC emergence (**Figure 6E**). Consistent with the reduced inflammatory profile observed in the hematopoietic tissue, baseline *il1β* expression was significantly decreased in tailfins from ALK5-inhibited animals (**Figure 6F**), indicating that embryonic ALK5 inhibition establishes a long-term dampened inflammatory state in peripheral tissues as well. Surprisingly, tailfins from ALK5-inhibited animals displayed elevated basal *mpeg1.1* expression as compared with DMSO controls (**Figure 6F**), suggesting increased macrophage accumulation under homeostatic conditions in peripheral tissues. Together with the reduced macrophage abundance detected in the WKM (**Figure 6B,C**), these findings suggest a systemic redistribution of macrophages from hematopoietic compartments toward peripheral tissues, accompanied by decreased inflammation.

The cytokine IL1β is a key mediator of regenerative inflammation for which macrophages represent a major cellular source during wound healing [48]. Therefore, we assessed tailfin wound regeneration in adult zebrafish exposed during embryogenesis to either DMSO or the ALK5 inhibitor. We investigated the expression of both *il1β* and *mpeg1.1* following injury. We observed an increase of *il1β* expression following injury in both conditions, with the ALK5-inhibited condition displaying expression levels comparable to DMSO controls (**Figure 6F**), indicating that *il1β*-inflammatory activation can still be induced upon tissue damage, despite impaired basal inflammatory conditions. However, overall macrophage abundance within wounded tailfins, assessed by *mpeg1.1* expression, was not significantly different between DMSO- and ALK5-treated groups (**Figure 6F**), raising the possibility that the observed strong inflammatory response may not be explained by changes in macrophage abundance, but rather by altered macrophage inflammatory activity or contributions from different cellular sources.

Tailfin regeneration was analyzed at 12 days post amputation (dpa), a stage at which macrophages constitute the predominant immune population within regenerating tissue [40]. The majority of ALK5-inhibited fish exhibited an altered pattern of tail regeneration, specifically in the dorsal tailfin region (**Figure 6G**). In contrast, the regrowth timing and tail length were similar between the two conditions (**Figure S3C**). Consistent with our findings, a previous study in adult zebrafish demonstrated that macrophage ablation during tissue regeneration resulted in fin patterning defects rather than impaired growth [49]. We therefore examined macrophage localization within regenerating tissues and observed a marked enrichment of macrophages in the dorsal-proximal region of embryonic ALK5-inhibited tailfins, corresponding to the area displaying defective regeneration (**Figure 6H**). This finding is consistent with previous evidence demonstrating that transient, but not sustained inflammatory signaling was needed for proper tissue regeneration [50]. The combination of altered basal inflammatory tone, systemic macrophage redistribution, and defective regenerative patterning supports the idea that developmental TGF-β signaling establishes long-term programs controlling macrophage homeostasis and inflammatory behavior in peripheral tissues and during tissue repair.

## DISCUSSION

In this study, we uncovered how transient embryonic cues during a critical developmental window exert durable effects on the hematopoietic system. Specifically, we illustrated that embryonic ALK5/TGF-βRI signaling during HSPC specification during ontogeny regulates HSPC dynamics within developmental hematopoietic niches, shifts lineage priming and macrophage differentiation programs, and imposes long-term effects on hematopoietic clonal architecture and inflammatory states in adult hematopoietic and peripheral tissues. Our findings indicate that signals acting within a critical developmental window impart properties on newly emerging HSPCs that are not corrected by subsequent developmental cues. Thus, even a temporary disruption of developmental signaling may permanently alter the trajectory of stem cell behavior and lineage specification thereby influencing susceptibility to hematologic disorders later in life.

Despite being embryonically-confined transient environments, developmental hematopoietic niches ensure the correct acquisition of both identity and architecture of the future hematopoietic and immune systems. Several signaling cues from the hematopoietic compartments are crucial for the correct development of the hematopoietic system [6–9, 51, 52]. Among them, TGF-β signaling was previously described as a key regulator of HSPC emergence, programming the vascular endothelium within the dorsal aorta to become hemogenic endothelium in collaboration with the transcription factor RUNX1 [14, 15]. The dorsal aorta is the primary embryonic vascular niche where definitive HSPCs are generated through an endothelial-to-hematopoietic transition. Nascent HSPCs subsequently migrate from the dorsal aorta to secondary niches, such as the CHT in zebrafish or the fetal liver in mammals. Here, they interface with resident endothelial cells, stromal cells, and primitive myeloid cells that aid in HSPC maturation, expansion, and differentiation [34]. By integrating live imaging, single-cell tracking, and transcriptional profiling, we identified that embryonic TGF-β signaling regulates early HSPC dynamics within the dorsal aorta that promotes egress and colonization into the CHT and regulates expression of migration-associated and lineage-specific genes (**Figures 1C,D and 3D,E**). Consistent with our findings, prior studies in mammalian HSPC emergence linked RUNX1 and TGF-β signaling to the regulation of numerous genes involved in hematopoietic lineage, adhesion and migration [15], suggesting one potential molecular mechanism for our observed ALK5 inhibitor effects on dorsal aorta HSPC dynamics. Moreover, a recent study also linked TGF-β signaling within the dorsal aorta to HSC migration to and maturation within the murine fetal liver, implying conservation for the role of this pathway and process in mammals [53]. Because migration into supportive hematopoietic niches is tightly coupled to lineage specification and differentiation [25, 54], our results imply that signaling based regulation of dorsal aorta HSPC delamination and appropriately timed CHT colonization is an early developmental bottleneck that influences lineage output and long-term hematopoietic composition.

While prior studies showed that TGF-β1 levels preferentially promoted proliferation and differentiation of adult myeloid-biased over lymphoid-biased HSCs [11, 16], they did not discriminate differentiation outcomes among specific myeloid cell fates. Building on these findings, we observed fewer macrophages in ALK5-inhibited larvae (**Figure 4D,F**) with no alterations in neutrophils (**Figure S1**), suggesting cell-type selectivity in TGF-β signaling regulation of myeloid cell fate outputs. The mechanisms underlying the selective impairment of the macrophage compartment, while sparing other myeloid-derived lineages, remain to be fully elucidated but the connection to altered niche dynamics hint that differences in cytokine availability and/or spatial signaling could underlie the immune cell-type selectivity.

Beyond a decrease in macrophage number, ALK5 inhibition seemed to shift the remaining macrophages toward a less inflammatory, more M2-like phenotype (**Figure 5C-G**), indicating that embryonic signals not only influence immune cell abundance but also preconfigure their functional competences. In certain contexts, like tumor microenvironments, TGF-β has been associated with the suppression of pro-inflammatory cytokine production and shifting macrophages toward an M2 phenotype [37, 39]. Our results highlight potential differences of TGF-β signaling on shaping differentiation of developing HSPCs versus the impact on adult macrophages. As it has been shown that TGF-β-mediated effects can be modulated by inflammation [55, 56], it is possible that the high sterile inflammatory milieu in the embryonic hematopoietic environments imposes a more pro-inflammatory TGF-β response in nascent HSPCs leading to the observed differences on macrophage subtypes compared to adult settings.

A major finding of the study was that embryonic TGF-β perturbation had lasting impacts on hematopoietic features known to depend on inflammatory inputs. Prior work demonstrated that inflammatory states influence the competitive fitness of hematopoietic clones, particularly within myeloid-biased populations. Elevated inflammatory cytokines, including IL-6 and IL-1β, can promote clonal hematopoiesis [57, 58]. In this context, macrophages can sustain pro-inflammatory states by expressing high levels of these pro-inflammatory cytokines, thereby reinforcing clonal selection [59, 60]. Paradoxically, in clonal hematopoiesis, dominant mutant HSCs often show a comparatively attenuated inflammatory transcriptional response relative to non-dominant or wild-type counterparts, suggesting that resistance to inflammatory stress may contribute to clonal selection [41, 61, 62]. In line with this logic, we demonstrate that transient embryonic ALK5 inhibition reduced inflammatory gene expression and the abundance of pro-inflammatory macrophages in the adult marrow, while promoting myeloid clonal selection (**Figure 6B-D**). As this shift in clonal composition occurred in the absence of additional mutations, it suggests that the inflammatory state alone might influence the competitive fitness of distinct HSPC subsets through non-genetic mechanisms. Moreover, it introduces the possibility that environmental cues encountered during development may represent previously underappreciated determinants of clonal hematopoiesis susceptibility. In support of this model, a prior study illustrated that embryonic HSPC-niche cell interplay modified adult hematopoietic clonality [63]. Combined, these findings identify embryonic developmental programming, through cellular and signaling niche components, as significant newly uncovered determinants of long lasting hematopoietic clonal selection.

The consequences of embryonic TGF-β perturbation extended beyond hematopoietic tissues into peripheral inflammatory processes. Tissue regeneration requires a tightly coordinated inflammatory response in which the spatial and temporal regulation of inflammatory signals critically determines repair outcomes [43, 64]. Although inflammatory responsiveness remained inducible in embryonic TGF-β-inhibited animals following tail fin injury (**Figure 6F**), the altered distribution of macrophages within regenerating tissues (**Figure 6H**) together with defective regenerative patterning (**Figure 6G**) suggests that the coordination and localization of inflammatory activity might be disrupted. As embryonic TGF-β inhibition does not abolish inflammatory responsiveness, it suggests that developmental imprinting primarily affects cellular identity and inflammatory organization rather than the intrinsic ability of cells to respond to wound stimuli. Clonal hematopoiesis is associated with altered differentiation, tissue distribution, inflammatory phenotype, and function of peripheral tissue macrophages [65], so it is attractive to posit that the inflammatory shifts in the tail fin reflect preferential infiltration by less inflammatory monocytes/macrophages originating from the clonally-shifted marrow compartment. However, we cannot exclude a contribution from embryonically-derived tissue-resident macrophages, whose developmental programming may have been altered also by embryonic TGF-β inhibition.

In summary, our study reveals that transient developmental perturbations can durably reshape hematopoietic organization, with consequences on hematopoietic imbalance and inflammatory dysfunction later in life. We identified embryonic TGF-β signaling as a critical developmental regulator of HSPC dynamics within their ontogenic niche that reshaped lineage output and inflammatory programming, imposing persistent effects on hematopoietic and inflammatory tone and clonal architecture. These findings add a new layer for understanding risk factors for adult clonal hematologic and immune disorders.

## MATERIALS AND METHODS

### Animal models

#### Zebrafish

Zebrafish were maintained according to Institutional Animal Care and Use Committee (IACUC)-approved protocols (#1636) in accordance with the Albert Einstein College of Medicine research guidelines. Wild-type zebrafish AB and transgenic lines were used in this study. Transgenic zebrafish lines used in this study included *Zebrabow-M* [66], *draculin:CreERT2* [67], *mpeg1.1:egfp* [68] and *mpeg1.1:mCherry* [69], *Tg(lyz:NTR-mCherry*) [70], *gata2b:Gal4;UAS:lifeactGFP* [71], *tnfa:egfp* [72], *Tg(gata1a:dsred*) [73], *runx1+23:mCherry* [32].

Embryos were maintained in embryo E3 medium [5mM NaCl (Fisher Scientific), 0.17 mM KCl (Dot Scientific), 0.33 mM CaCl2 (Acros Organics), 0.33 mM MgSO4•7x H2O (Sigma-Aldrich)] until 5dpf. Thereafter, they were moved into the Fish Facility system.

#### Drug treatment

TGF-β receptor I (ALK5) was inhibited by treatment with the SB431542 (Selekchem-S1067) drug at a concentration of 20 μM. Embryos were first dechorionated. ALK5 inhibitor or DMSO were added to embryo E3 medium in 6-well plates with 40-50 30hpf-embryos per well. Treatment was stopped at 50hpf by 3 washes in embryo E3 medium. 50hpf embryos were then moved to petri dishes.

#### Cell collection and flow cytometry

20-30 embryos were anesthetized with 0.01% tricaine (Fisher) and then resuspended in a Dissociation buffer [HBSS (Life Technologies) containing D(+) Glucose (Sigma), Fetal Bovine Serum (ThermoFisher), Collagenase Type IV (ThermoFisher)] at 30*C for 20-30min [74]. A single-cell suspension was mechanically prepared by pipetting up and down vigorously. The enzyme reaction was then stopped with the addition of cold FACS buffer (0.9x D-PBS, 5% FBS, 1% Penn/Strep (Life Technologies)) and pelleted.

Adult fish (3- and 6-months post fertilization) were anaesthetized with 0.02% tricaine in embryo E3 medium and dissected to collect the WKM in FACS buffer. A single-cell suspension was mechanically prepared by pipetting up and down vigorously. The lysis of red blood cells (RBCs) was done with an ammonium chloride-based osmotic shock lysis protocol, previously described in [75]. Briefly, cells were resuspended for 10 min reaction within an RBC Lysis buffer (0.17 M Tris-HCl pH 7.65 and 0.16 M NH4Cl (1:10)), the reaction was stopped by cold FACs buffer.

Embryo, larval and adult zebrafish cell suspensions were filtered through a 40μm cell strainer (Falcon), pelleted by centrifugation, and resuspended in 200 μL of FACs buffer containing either 1 μg/mL of DAPI (Sigma D8417) or 2 nM DRAQ-7 dead cell dye aka Draq7 (Invitrogen, D15105).

Data were acquired at the Flow Cytometry Core Facility at the Albert Einstein College of Medicine using an Aurora [5L/UV/V/B/YG/R] (BD Biosciences) and were then processed using FlowJo Software (versions 10.6 - 10.7.1).

#### Whole mount *in situ* hybridization and imaging analyses

After treatment with DMSO or ALK5 inhibitor, 50hpf embryos were fixed in cold 4% PFA and stored at 4°C until usage for WISH. Whole mount in situ RNA hybridization was done based on established zebrafish protocols [76]. Pigment removal by bleaching, Proteinase K treatment and 0.25% glutaraldehyde fixation were each done for 20 minutes at room temperature. To visualize RNA probe signal, we used anti-DIG followed by colorimetric detection using NBT/BCIP. Samples were incubated in NBT/BCIP for 4 hours at room temperature. The following probe was used: cmyb [9]. TIF images were processed in FiJi [version 2.1.0/1.53c]. Integrated density was derived by subtracting the background signal and normalizing to the area of the Region of interest (ROI). Data were analyzed in Excel and Prism.

### Fluorescent imaging

Live zebrafish embryos (50 hpf – 6 dpf) were anesthetized with 0.01% tricaine (Fisher), then mounted in 4% (wt/vol) methylcellulose in 35-mm imaging dishes (MatTek) as described previously [77]. Fluorescent imaging of transgenic fluorescent zebrafish embryos was performed with Zeiss Discovery.V8 and Zeiss Axio Observer A1 Inverted microscope with an AxioCam HRc Zeiss camera and Zeiss Zen 2 or 2.5 software. Fluorescence was detected with cyan fluorescent protein (CFP), mCherry, Texas Red (for dsRed lines), and green fluorescent protein (GFP) filters.

Fluorescence microscopy images were analyzed using FIJI (Schindelin et al., 2012). On each image, a region of interest (the fluorescent tissue within the CHT or thymus) was manually drawn. Corrected Total Cell Fluorescence (CTCF) was calculated as Integrated density of region of interest - (Area of region of interest X Mean fluorescence of 4 different background regions).

### Live imaging of HSPC migration, and data processing and quantitative analysis of HSPC migratory behavior

Live imaging of HSPC dynamics was performed using *Tg(gata2b:Gal4;UAS:lifeactGFP*) embryos. 50hpf embryos were anesthetized and loaded into a zWEDGI chamber for time-lapse imaging, as previously described [78]. LMP agarose (2%) (Sigma-Aldrich) in tricaine/embryo E3 medium (∼28°C) was placed over the larva’s head and then allowed to solidify with the embryo in the proper position. Additional tricaine/embryo E3 medium was added to the chamber as needed. Live imaging of embryos was assessed using a spinning-disc confocal microscope Nikon CSU-X; Yokogawa. 2 × 1 tile images were taken and automatically stitched. Time-lapse was recorded every 2min for 20 hours, 20x air objective. All images were processed using IMARIS Bitplane software (Version 9.5/9.6). Number of HSPCs was automatically counted in the whole embryos using IMARIS spots function. Spots were defined as particles with 5 μm and 10 μm of X/Y and Z diameter, respectively. Tracking data obtained from IMARIS were exported as merged tables containing positional, displacement, velocity and track-based features for each HSPC over time. Downstream analyses were performed in R (version 4.5.2). To ensure data quality, duplicated entries were identified based on TrackID, pID.x, TimePoint and cell ID, and resolved by retaining the entry with the lowest number of missing values. Rows containing more than 50% missing values were excluded from further analysis. Datasets from independent experiments and biological replicates were annotated with condition (DMSO or ALK5 inhibitor), tissue (DA or CHT), and replicate identifiers, and subsequently merged into a unified dataset. Features associated with imaging artifacts or redundant measurements (e.g., intensity-related variables or axis-specific components) were excluded prior to analysis. To ensure temporal comparability across experiments, timepoints were rescaled for each replicate to a common time window (0–30,000 seconds), allowing alignment of migration dynamics across embryos. For time-course analyses, the number of tracked HSPCs per timepoint was calculated and normalized to the initial timepoint (t0) for each replicate.

For feature-based analyses, quantitative parameters describing cell migration (e.g., displacement length, track duration, speed and neighborhood distance metrics) were extracted and standardized using z-score normalization across all cells. This normalization enabled comparison of relative feature values between conditions.

Dimensionality reduction was performed using Uniform Manifold Approximation and Projection (UMAP) on the selected migration features to capture HSPC behavioral states. Clustering of single-cell tracks was performed based on these features to define distinct behavioral clusters within each tissue.

For comparative analyses between DMSO and ALK5 inhibitor-treated conditions, statistical differences in feature distributions were assessed using two-tailed Mann–Whitney tests. Multiple testing correction was applied using the Benjamini–Hochberg method when appropriate.

### scRNA-sequencing and data processing

Flow cytometry protocol was used to prepare *Tg(mpeg1.1:mCherry)* (6 dpf) and *Tg(gata2b:Gal4;UAS:lifeactGFP)* (3dpf) for fluorescent cell sorting prior to 10X Genomics scRNA-seq. Per sample, 200-300 larvae were used. GFP^+^ cells were sorted at 3dpf, mCherry^+^ cells were sorted at 6dpf on BD FACSAria II Cell Sorter, equipped with a 100-micron nozzle. Sorted cells were collected into microcentrifuge tubes containing 500 μL IMDM + 10% FBS + 1% Pen/Strep at 4°C and processed for library preparation using the 10X Genomics Chromium GEM-X Single Cell 3ʹ Kit v4.1 PN-1000691, performed by the Einstein Genetics Genomics Core. Libraries were then sent to Genewiz for sequencing.

After sequencing, data processing was assessed as previously described [79]. Briefly, Fastq files were processed in Cell Ranger 8.0.1 using the counts function to generate the matrix. A custom reference genome was built based on *Danio rerio* GRCz11 with the addition of mCherry, GFP, and T cell specific genes: TCRaC, TCRbC1 and TCRbC2. All subsequent dimension reduction and clustering was done in Seurat version 5.2.0. [80]. Cells with less than 200 genes and more than 10% mitochondrial DNA were filtered out.

For the 3 & 6 dpf datasets: initial clustering was done using 4,000 top variable features, 20 principal components (PC) and at cluster resolution of 0.5. No batch effect was apparent between DMSO and ALK5 inhibited conditions, so analysis was done without batch corrections. The 3 dpf dataset contained non-hematopoietic populations like melanocytes, neurons, and hepatocytes, so the cells were filtered out to contain only HSPC populations on the basis of *myb* and *EYFP* gene expression. The final HSPC subset was clustered using 4000 top variable features, 20 PCs, and 0.1 cluster resolution. The 6 dpf dataset contained non-hematopoietic lineage cells like neurons, mesenchyme cells, and epidermis so cells were filtered out to keep only macrophages expressing *mpeg1.1* and *mCherry* markers. From the initial 20 clusters, cells were further filtered and clusters 0, 3, 8, 11, 15, 16 were kept and clustered using 20 PCs and a resolution of 0.2. The final clustering was obtained by keeping clusters 0, 1, 3, 5, 6 on the basis of *mfap4* and *mpeg1.1* expression. Final clustering of 6 dpf macrophages was done using 20 PCs and a cluster resolution of 0.2.

For cluster identification, we used a combination of literature-based marker genes and an assessment of top markers using the standard FindAllMarkers function in Seurat which uses the Wilcoxon Rank Sum test.

For differential gene expression analysis, the DMSO and ALK5 inhibited conditions were compared using the FindMarkers function in Seurat using standard settings for Wilcoxon Rank Sum test. The GSEA analysis was done using the gseGO function in ClusterProfiler [81]. The differentially expressed genes were ordered by fold-change and subject to GSEA analysis using the Biological Process gene ontology category. Selected terms are plotted, see Supplemental files 1 and 2 for full list of resulting terms. Module scoring for M1/M2 signature genes was done using the built in AddModuleScore() function in Seurat. See Supplemental File 2 for list of genes used in each module.

All scRNA-seq data are available via the GEO public repository under accession number GSE341887.

### RT-qPCR

Zebrafish whole embryo RNA was isolated from dechorionated embryos using the combined Trizol (Invitrogen) and PureLink RNA Mini Kit (Invitrogen) protocol according to the manufacturer’s instructions. RNA from adult fish WKM was extracted using the Quick-RNA Microprep Kit (Zymo Research) TURBO DNA-free kit (Life Technologies) was used for DNA removal following RNA extraction. Isolated RNA was quantified using Qubit RNA High Sensitivity Kit (ThermoFisher) and stored at −80°C. cDNA was synthesized using the High-Capacity cDNA Reverse Transcription Kit with RNase Inhibitor (Life Technologies) according to the manufacturer’s instructions. The expression levels of cellular markers and inflammatory cytokines were evaluated using qPCR [Fisher QuantStudio™ 6 Pro Real-Time PCR System] and the SYBR Green kit (ThermoFisher). Relative mRNA expression levels were obtained by standardizing to the *rps11* mRNA level using the 2−ΔΔCt method. The specific primers used in this study are listed in **Supplementary Table 1**.

### Zebrabow

To study shifts in clonality in zebrafish, we used a system called Zebrabow [66]. Briefly, the Zebrabow encodes for a transgene that has three fluorophores (dTomato, mCerulean and eYFP) in tandem. Cells can have up to 20 copies of this transgene, and, in the inactivated state, they express only the dTomato fluorophore. Upon a tamoxifen treatment, the Cre-mediated stochastic recombination of the lox sites between the three fluorophores leads to the expression of the mCerulean and eYFP fluorophores as well. As cells carry different zebrabow transgene copy numbers, each cell will have a unique color barcode and its progeny will inherit this barcode throughout life, allowing the study of clonal evolution. Optimized Zebrabow labelling using *draculin:creERT2* allows the assessment of the clonal architecture of the hematopoietic compartment [42]. In our study, 24hpf *Tg(ubi:Zebrabow-M;draculin:CreERT2*) embryos were treated with 15 μM 4-hydroxytamoxifen for 6 hours in the dark at 28.5°C, then placed in fresh embryo E3 medium. Embryos with dim transgene expression were excluded for analysis to account for variation in Zebrabow transgene insert number. WKM from adult zebrafish (older than 3 months) was isolated and processed for flow cytometry acquisition as described above. At least 10,000 events were acquired in the FSC/SSC myelomonocyte gate to be used for later clustering. FlowJo™ v10 Software was used to gate myeloid, precursors, and lymphoid populations based on FSC/SSC characteristics. Additionally, cells that had no detectable CFP and YFP expression (TdTomato-only) were discarded in the analysis because they derive from stem cells that did not undergo Zebrabow-recombination. Zebrabow cluster analyses, including the identification and quantification of the diverse cluster per sample and their own size, were assessed using previously published pipelines adapted to a Python-based interface [41, 42]. Only zebrafish WKM samples with greater than 75% recombination efficiency were included into analyses.

### Tailfin injury and regeneration phenotype assessment

Fin regeneration was assessed in adult (3-9 months old) *Tg(mpeg1.1:egfp*) zebrafish treated during embryogenesis with DMSO or ALK5 inhibitor as described above. Adult animals were anesthetized with 0.02% tricaine and the distal half of their caudal tailfin was amputated with a sterile razor blade. Imaging of fin regeneration and *mpeg1.1*:EGFP^+^ cell localization was performed with Zeiss Discovery.V8 and Zeiss Axio Observer A1 Inverted microscope with an AxioCam HRc Zeiss camera and Zeiss Zen 2 software. Tail fin area and egfp fluorescence intensity profile within the tail were assessed using the FiJi software. Fin regeneration appearance was classified in two categories, normal or abnormal in two independent experiments.

Caudal fin regrowth was measured in Fiji using the straight-line tool. For each fish, three measurements were taken in the dorsal, medial, and ventral regions of the regenerating fin (see Supplementary Figure 3C), and the lengths of these lines were used to evaluate fin regrowth. To assess the distribution of *mpeg1.1:eGFP^+^* cells within the tailfin, dorsal and ventral regions of the wounded fin were defined using the Fiji rectangle tool. The EGFP mean fluorescence intensity (MFI) was measured in each region, and the ratio of ventral-to-dorsal EGFP signal was calculated for each fish.

### Quantification and statistical analysis

Statistical analyses were carried out using GraphPad Prism V.10. Experiments were performed with a minimum of two independent replicates. Details are depicted in individual figure legends. The P values are represented as follows: ****P < 0.0001, ***P < 0.001, **P < 0.01, *P < 0.05, and not significant (ns) P ≥ 0.05.

## Data availability

All scRNA-seq data are available in the GEO NCBI public repository under accession number GSE341887.

## Acknowledgments

M.M. was supported by an American-Italian Cancer Foundation Post-Doctoral Research, a Paul Frenette Scholar Award from the Ruth L. and David S. Gottesman Institute for Stem Cell Research, a Ceriale postdoctoral fellowship from the Einstein-Montefiore Blood Cancer Institute, an American Cancer Society Postdoctoral Fellowship. T.V.B. was supported by grants from the National Institutes of Health (NIH) R01DK121738, R01DK131445, R01DK141169, R01DK142340, and the Edward P. Evans Foundation. A.N. was supported by NIH F31HL167600, NYSTEM award C34874GG (PI Frenette), and a Liang Zhu Memorial Fellowship. SDO was supported by NIH R35GM147416.

The authors would like to thank all members of the Bowman laboratory for meaningful discussions and technical assistance, and the Einstein Stem Cell Institute and Developmental and Molecular Department of Albert Einstein College of Medicine for their comments. The authors would like to thank the Einstein Zebrafish, Flow Cytometry, Analytical Imaging, and Genomics core facilities. Flow cytometry and scRNA-seq studies were carried out using resources of the FACS Core Facility and Genomics Facility of the Montefiore-Einstein Comprehensive Cancer Center, which is supported by NIH/NCI Cancer Center Service Grant P30 CA13330 and by Shared Instrumentation grants S10OD026833 and S10OD032169.

## Contributions

Conceptualization, experiment design: MM, TVB. Investigation: MM. Single cell data curation and analyses: MM, AN, TVB. Cell dynamic behavior methodology, data curation and analyses: MM, JCS, SDO. Original draft writing: MM, TVB. Review and editing of the manuscript: AN, JCS, SDO, MM, TVB.

## Declaration of Interests

The authors declare no competing interests.

## Supplementary Figures, Legends, and Tables

**Figure S1.**
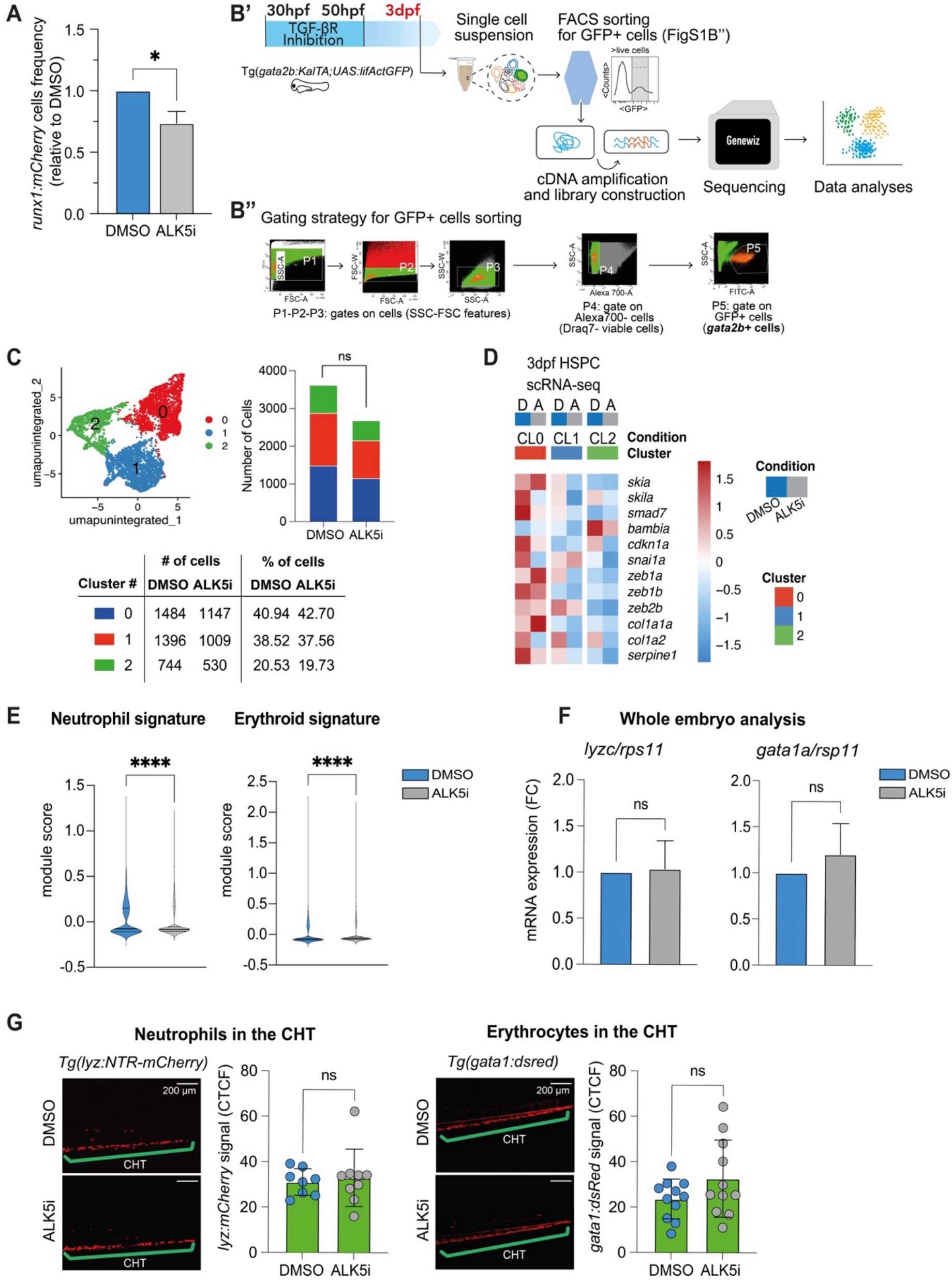
**A.** HSPC frequency in *Tg(runx1+23:mCherry)* larvae was quantified by flow cytometry at 3 dpf. Bar graph represents mean + SEM of 4 biological replicates. Statistical differences were calculated by Mann-Whitney unpaired test. *P < 0.1. **B’.** Experimental workflow for single-cell transcriptomic profiling of HSPCs from 3 dpf *Tg(gata2b:KalTA;UAS:lifActGFP*) larvae treated embryonically with DMSO or ALK5 inhibitor. **B’’.** Representative FACS gating strategy for isolation of viable GFP^+^ (*gata2b^+^*) cells. Sequential gates (P1–P3) were used to select cells based on forward and side scatter (FSC/SSC) properties. Dead cells were excluded using Draq7 staining (P4), and GFP^+^ cells (Alexa700⁻/GFP⁺) were selected in the final gate (P5) for sorting. **C.** Single-cell transcriptional analysis identified 3 clusters of HSPCs in both DMSO and ALK5-inhibition conditions. Graph on right shows the number of cells per cluster. Both DMSO and ALK5 inhibitor-treated samples contained a major cluster (Cluster 0), which was composed of 1484 cells in DMSO and 1147 cells in ALK5, followed by Cluster 1 composed of 1396 cells in DMSO and 1009 cells in ALK5, and a smaller cluster (Cluster 2) composed of 744 and 530 cells, respectively in DMSO and ALK5. Number of cells, as total number and percentage is reported in the table. **D.** Heatmap showing selected differentially expressed TGF-β signaling pathway and target gene profiles across the different clusters and conditions. Each row represents a unique gene. Each column represents the DMSO *vs* ALK5 inhibited conditions in each cluster. Relative gene expression levels are signified by different color scale (red=higher and blue=lower). **E.** Quantification of neutrophils- and erythroid-associated gene module scores in DMSO and ALK5 inhibitor conditions across all clusters. List of genes used for module score is in **Supplementary File 1**. Statistical differences were calculated by Wilcoxon test. ****P < 0.0001. **F.** RNA expression of *lyzc* and *gata1a* from 6 dpf larvae was measured by RT-qPCR. Values were normalized to *rps11* and expressed as fold change (FC) (3 independent experiments). **G**. *Tg(lyzc:NTR-mCherry)* neutrophils (left) and *Tg(gata1a:dsred)* erythrocytes (right) were quantified in the CHT of 6 dpf larvae. Representative fluorescent images are shown: DMSO (top) and ALK5 inhibited (bottom). Scale bars, 200 μm. Bar graphs show the Corrected Total Cell Fluorescence (CTCF) (number of larvae > 10). Statistical differences were calculated by Mann-Whitney unpaired test. ns, not significant.

**Figure S2.**
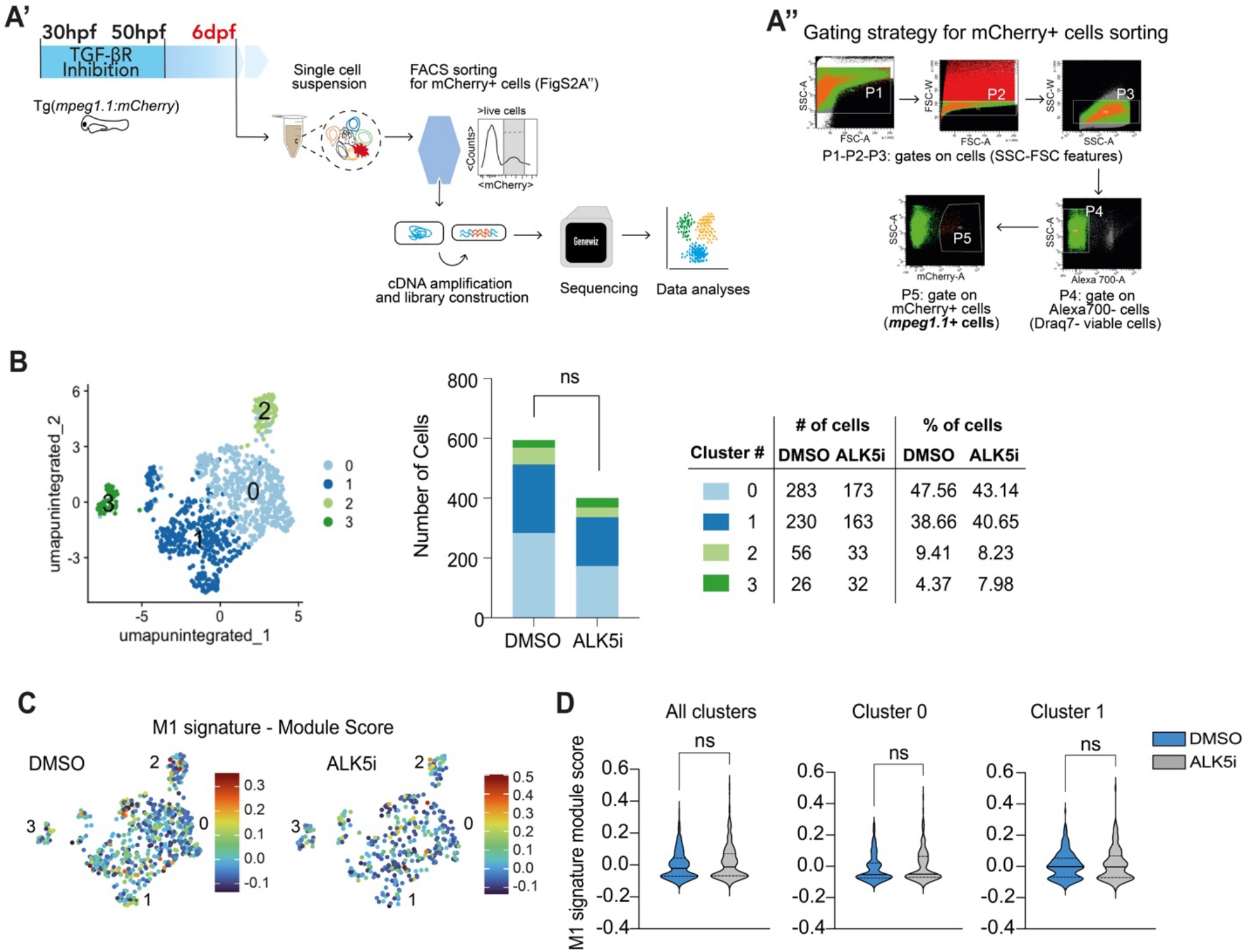
**A’**, Experimental workflow for single-cell transcriptomic profiling of macrophages from 6 dpf *Tg*(*mpeg1.1:mCherry*) larvae treated embryonically with DMSO or ALK5 inhibitor. **A’’.** Representative FACS gating strategy for isolation of viable mCherry^+^ (*mpeg1.1+)* cells. Sequential gates (P1–P3) were used to select cells based on forward and side scatter (FSC/SSC) properties. Dead cells were excluded using Draq7 staining (P4), and mCherry^+^ cells were selected in the final gate (P5) for sorting. **B.** Single-cell transcriptional analysis identified 4 clusters of macrophages in DMSO and ALK5-inhibition conditions. Graph showing the number of cells per cluster. Both DMSO and ALK5 inhibited samples presented two major clusters (Clusters 0 and 1), which were composed by 283, 230 and 173, 163 cells, respectively in DMSO and ALK5 inhibited conditions. Two minor clusters (Clusters 2 and 3), were composed by 56, 26, and 33, 32 cells, respectively in DMSO and ALK5 inhibited groups. Number of cells, as total number and percentage is reported in the table. **C,D**. Module scores for M1-associated genes in DMSO and ALK5-inhibited conditions with quantification across all clusters and within clusters 0 and 1. Full lists of signature genes are listed in **Supplementary File 2**. Color in the feature plots indicates the M1 macrophage module score, where positive values denote enrichment and negative values denote depletion of the M1-macrophage transcriptional program. Statistical differences were calculated by Wilcoxon test. ns, not significant.

**Figure S3.**
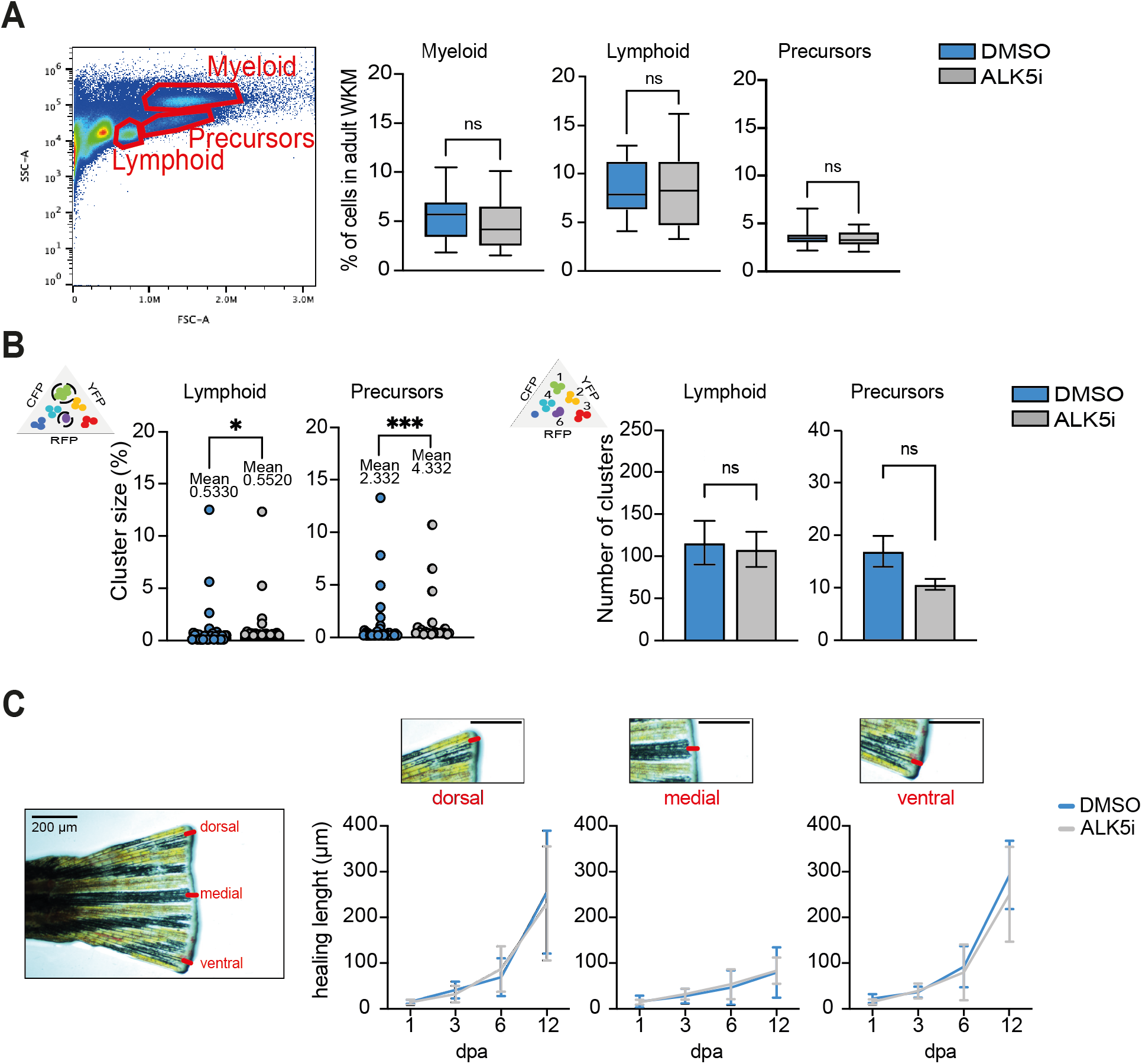
**A,B**. *Tg*(*ubi:Zebrabow-M;drl:creERT2*) embryos were treated with 4-OHT from 24-30hpf. They were then treated with DMSO or ALK5 inhibitor from 30-50 hpf and grown to adulthood. Whole Kidney Marrow (WKM) was extracted and single cells suspension was prepared for flow cytometry. Dead cells were excluded from the flow cytometry gating using Draq7 staining. Schematic of the experiment is in Figure 6D. The graphs to the right show the percentage of myeloid, lymphoid, and precursor cells quantified by flow cytometry in 2 independent experiments, N> 17 total animals per group. Statistical differences were calculated by Mann-Whitney unpaired test. ns, not significant. **B.** Flow cytometry data from lymphoid and precursor cells (defined by FSC/SSC parameters) were clustered using a custom MATLAB graphical tool developed Avagyan *et al.*, 2021 [41]. Cluster size and cluster number were analyzed in two independent experiments, N=16-20 animals per group. Statistical differences were calculated by Mann-Whitney unpaired test. *P < 0.05, ***P ≤ 0.001, ns, not significant. **C**. Fin amputation was assessed in adult zebrafish treated during development with DMSO or ALK5 inhibitor from 30-50hpf. Schematic of the experiment is in Figure 6E. The kinetics of wound healing was assessed at 1, 3, 6, and 12 days post amputation (dpa). A representative brightfield image of a regenerating zebrafish caudal fin illustrating the three anatomical regions used for regenerative measurements: dorsal, medial, and ventral. Red lines indicate the position at which regenerative outgrowth or “healing length” was measured. Quantification of healing length is plotted separately for dorsal, medial, and ventral fin regions, with magnified examples of the corresponding measured regions above each graph. Scale bars, 200 µm. Data are from N=6-7 animals. Statistical differences were calculated by Mann-Whitney unpaired test and not significant, not shown.

**Supplementary Movie 1: Spinning disk imaging of 3 days post fertilization larvae for behavioral analysis of HSPCs dynamics within the DA**. Representative image sequence of the Dorsal Aorta (DA) area from *Tg(gata2b:KalTA;UAS:lifActGFP)* larvae treated embryonically with DMSO or ALK5 inhibitor. The DA is the hematopoietic niche where HSPCs emerge. After specification from the hemogenic endothelium of the DA, HSPCs exit the niche and migrate to the caudal hematopoietic tissue (CHT), where differentiation occurs. In DMSO controls, HSPCs rapidly leave the field of view as they exit the DA niche and enter circulation. In ALK5 inhibitor– treated larvae, *gata2b^+^* cells are more numerous within the DA compared to DMSO controls, indicating accumulation within the hematopoietic niche. This observation is consistent with the increased track duration quantified in **Figure 2E**. In addition, live imaging show that *gata2b^+^* cells from ALK5 inhibitor–treated larvae exhibit increased cycles of membrane extension and retraction, consistent with an increased dynamic (e.g. displacement length and speed variation) but non-productive motility pattern as reflected by their lower mean speed analyzed in **Figure 2E**. These features are indicative of impaired migratory behavior that may hinder HSPC exit from the DA and subsequent colonization of the CHT. The kinetic parameters extracted from this time-lapse were used for the analyses shown in **Figure 2A-B and 2E**. Scale bar, 10µm.

**Supplementary Movie 2: Spinning disk imaging of 3 days post fertilization larvae for behavioral analysis of HSPCs dynamics within the CHT**. Representative image sequence of the CHT area of *Tg(gata2b:KalTA;UAS:lifActGFP)* larvae treated embryonically with DMSO or ALK5 inhibitor. No obvious difference in the number of *gata2b^+^* cells within the CHT is apparent between DMSO- and ALK5 inhibitor-treated larvae. As shown in **Figure 2C–D**, the HSPC population in the CHT displays heterogeneous migratory behavior, comprising both relatively stationary cells and a more motile subset, that seems being predominantly located in the central region of the CHT. Notably, these highly motile cells exhibit faster migration in DMSO-treated larvae than in ALK5 inhibitor-treated larvae, consistent with the higher mean speed quantified in **Figure 2F**. Kinetics features of GFP^+^ cells were extracted and used for the analyses shown in **Figure 2C-D and 2F**. Scale bar, 10µm.

**Table S1:**
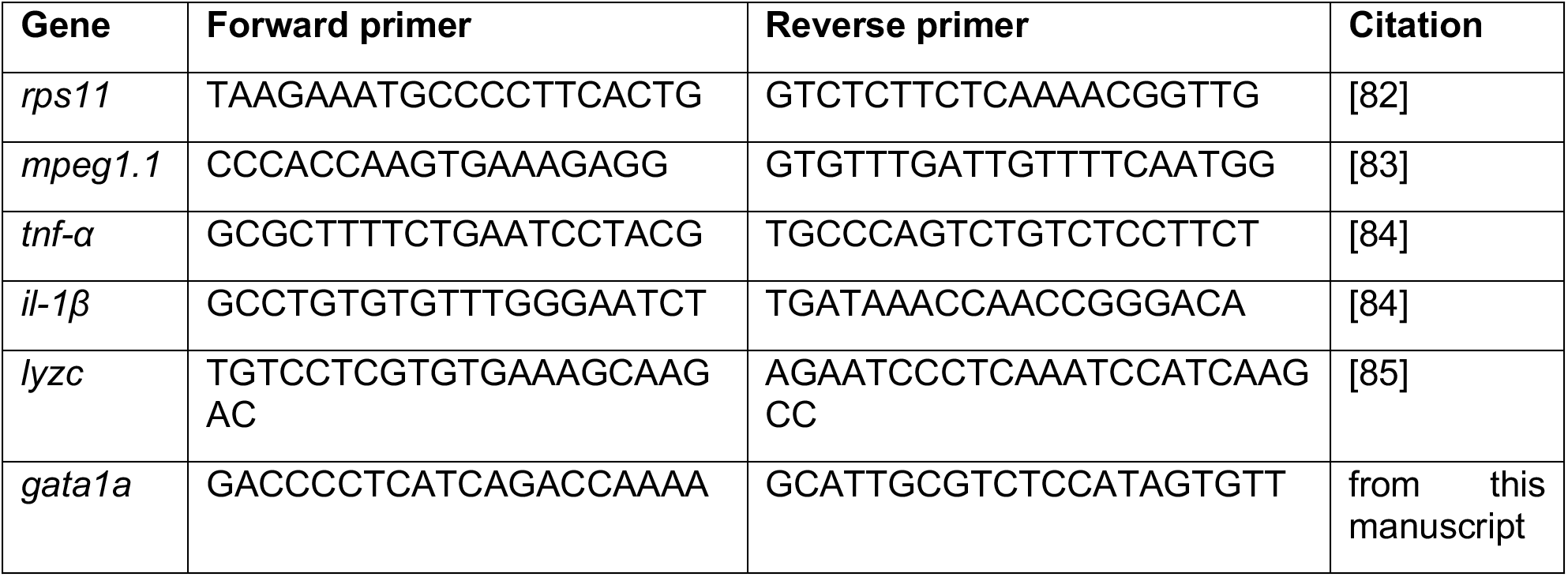
qPCR primers used throughout the paper.

## Notes

### Competing Interest Statement

The authors have declared no competing interest.

